# A New Frontier in CWD Detection: Antemortem Plasma Biomarkers and Behavioral Profiling in Transgenic Mouse Models

**DOI:** 10.64898/2026.08.04.742870

**Authors:** Alyssa L. Seerley Nolan, Serena D. McElroy, Allison A. Mace, Andrea L. Grindeland Panter

**Affiliations:** Weissman Hood Institute at Touro University; McLaughlin Research Institute Great Falls, MT 59405

## Abstract

Chronic Wasting Disease (CWD) is a fatal transmissible spongiform encephalopathy (TSE) that is confined to cervids (deer, moose, elk, and reindeer) but shares key properties with human neurodegenerative conditions such as Alzheimer’s, Parkinson’s, Huntington’s disease and frontal-temporal dementia. CWD and other TSEs are caused by the misfolded prion protein (PrP). Although the identification of diagnostic and prognostic biomarkers at all stages of disease progression is becoming exceedingly critical as CWD continues to increase in prevalence, accurate antemortem testing techniques are extremely limited. This study made use of cervidized transgenic mice (mice carrying the cervid PrP) that recapitulate CWD in various disease stages and investigated the utility of neurological biomarkers and neurobehavioral manifestations for CWD detection. Neurofilament light chain (NFL), glial fibrillary acidic protein (GFAP), and total Tau (t-Tau) were assessed under the hypothesis that combined biomarker signatures might more reliably reflect CWD-related neurodegeneration and disease progression. Analyses at 90, 132, 174, and 230 days post-CWD inoculation show distinct biomarker elevation, with all three biomarkers significantly elevated in the CWD animals by 132 days post-inoculation. To our knowledge, this is the first demonstration that these three plasma biomarkers are useful not only for detecting CWD, but also for identifying it at early antemortem stages of disease. Novel phenotypes were also revealed by comprehensive phenotypic profiling, including rigid tail elevation, increased grip strength, and impaired coordination, to lend further support to plasma biomarker data indicating neurologic impairment associated with brain pathology. Ultimately, the goal is to improve antemortem, non-invasive CWD detection methods to enable earlier detection and assist with disease management.

## 1. Introduction

Transmissible spongiform encephalopathies (TSEs), or prion diseases, are relatively rare but fatal neurodegenerative diseases that occur in a wide range of mammals and are characterized by a neuropathologic spongiform appearance. TSEs include, but are not limited to humans (Creutzfeldt-Jakob, fatal familial insomnia, Gerstmann-Sträussler-Scheinker), cattle (bovine spongiform encephalopathy), sheep (scrapie), and cervids (chronic wasting disease) ^1,2^. The central molecular mechanism across all TSEs is conserved in that the cellular form of the prion protein (PrP^C^) misfolds to the diseased-associated isoform (PrP^sc^) ^2–4^. TSEs often lead to neuronal death and vacuolization in the central nervous system that is hallmarked by an abundance of astroglial cells but is deficient in typical immune responses^5,6^. Despite extensive studies, knowledge regarding the physiological role of PrP^C^, the conversion to PrP^sc^, and the biochemical properties that impact disease etiology, transmission, and prevalence, remains limited.

CWD was first identified in a captive mule deer in Colorado in 1967 and has been spreading in both wild and captive animal populations ever since^7,8^. According to the United States Geological Survey (USGS), CWD was found in 37 states and 5 Canadian provinces in 2026. Unlike other prion diseases, CWD is highly infectious within cervid populations; CWD can be shed through peripheral systems including excrement and bodily fluids such as urine, saliva, blood, and antler velvet before clinical signs are displayed. Evidence indicates that the environment plays a major role in horizontal transmission as a result of the binding and uptake of infectious PrP^sc^ by plants, soils, and water, effectively contributing to the spread of disease in livestock and wildlife for years after initial shedding of prions ^9–12^.

Practical testing modalities for CWD are crucial for reducing disease spread. Current diagnostic methods remain largely laboratory-based, including ELISA, immunohistochemistry (IHC), and Real-Time Quaking-Induced Conversion (RT-QuIC), all of which require specialized reagents, equipment, and technical expertise for accurate interpretation. Most commonly, these assays rely on postmortem tissues, underscoring the need for more accessible sample types that support reliable antemortem testing. Although RT-QuIC has improved prion detection sensitivity, blood-based RT-QuIC remains challenging and has not yet become a practical diagnostic approach^13–15^. Field-deployable testing options are also extremely limited. Therefore, the development of less invasive assays using accessible tissues or fluids, such as blood, could substantially improve CWD surveillance and management.

In this study, we sought to overcome the limitations of working with wild animal populations, and the limited accessibility to samples infected with CWD, by employing a transgenic mouse model in our studies. Transgenic CWD models offer numerous advantages including the ability to assign research groups with statistically appropriate numbers and the ability to define onset of disease through timed inoculation. These advantages enable us to not only validate biomarkers for diagnosing disease but also determine the stage (early, middle, or late). Indeed, we have found that the novel combination of highly sensitive plasma biomarkers, including neurofilament light (NfL), glial fibrillary acidic protein (GFAP), and total Tau (t-Tau), allows continual live monitoring of disease progression in CWD transgenic mouse models that express wildtype cervid prion protein gene (*PRNP*). Furthermore, identification and characterization of the earliest, oftentimes subtle, neuropathological timepoints through phenotypic analysis identified improve live-animal diagnostic testing targets and can be used to evaluate therapeutic intervention when available^16^. Although behavioral phenotypic analysis of other neurodegenerative diseases has been viewed as a vital tool to track disease progression^16–18^, there are few behavioral studies published regarding CWD in mouse models, and those are limited in scope^19,20^.

Here, we report establishment of a system for phenotyping CWD mouse models that we have termed “comprehensive phenotypic profiling” (CPP). By integrating three primary plasma biomarkers with behavioral manifestations across categories such as observable traits, strength, coordination, and gait analyses, we can identify and categorize disease phenotypes across a spectrum of antemortem stages. Thus, here, we report our findings using the Tg1536 model which express wildtype cervid *PRNP*^1,2,21^.

## 2. Methods

### 2.1 Mice

All animal procedures were conducted in compliance with the guidelines outlined in the US National Research Council’s *Guide for the Care and Use of Laboratory Animals* ^22^ and the US Public Health Service’s *Policy on Humane Care and Use of Laboratory Animals*. The study protocol, AG-100, was reviewed and approved by the Institutional Animal Care and Use Committee (IACUC) at the McLaughlin Research Institute. Mice were housed in the Animal Resource Center at the Weissman Hood Institute at Touro, McLaughlin Research Institute, a facility exclusively for mice and accredited by the Association for Assessment and Accreditation of Laboratory Animal Care International.

This study primarily used the Tg1536 strain of mice which express cervid *PRNP* at a five-fold higher level than the level of wildtype PrP expression in FVB mice^1,2,21^. The strain was maintained with two copies of the transgenic allele on a mouse *PRNP* null, FVB/NJ background. To induce CWD (or produce appropriate controls), 7-week-old Tg1536 mice were intracerebrally inoculated in a biosafety level 2+ laboratory (BSL2+). A cohort of 20 mice were assigned to be in a group destined for all longitudinal experiments. Twelve mice (7 male, 5 female) were inoculated with CWD, and eight mice (4 male, 4 female) were inoculated with normal non-diseased mule deer brain homogenate (NBH). An additional cohort of 90 days post inoculation (dpi), 160dpi, and 230dpi (six CWD, four NBH) timepoint mice were inoculated and harvested for histology and biomarker purposes. Various control mice were also included. Tissues from one *PRNP* null mouse was used in RT-QuIC as a “no seeding” control. Ten FVB/NJ mice served as mouse *PRNP* wildtype background strain controls. Five were inoculated with CWD while the other five were inoculated with NBH. Five uninoculated Tg1536 mice and five uninoculated FVB mice were also used as controls, primarily in the biomarker assays. Four 5xFAD^23^ mice were used as an additional control exhibiting neurodegenerative disease. Table 1 provides a summary of the mouse model, the available number of mice per group, and the study they were utilized in. Each study method did not necessarily use all the mice available.

**Table 1:**
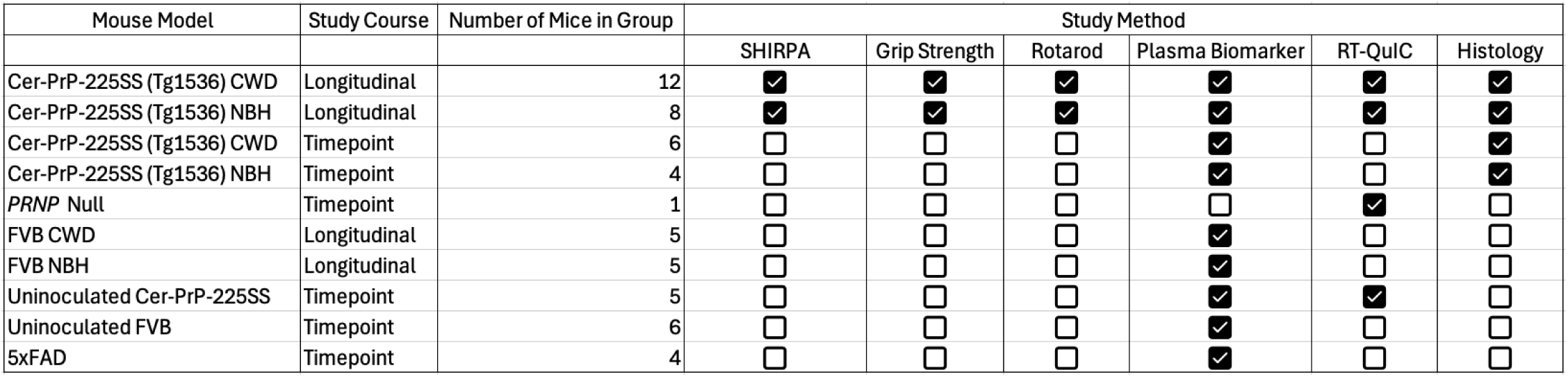
Breakdown of mice used in the study.

All inoculated mice were injected with 30uL of 1% CWD infected mule deer brain homogenate or NBH (*PRNP* wildtype) into the right thalamus using a SafetyGlide Insulin needle (31 x 6mm gauge) (VWR, #10002-698) under isoflurane anesthesia. After inoculations, meloxicam (MWI, #501080) was administered, and the mice were placed on warmers and monitored until deemed safe and alert. Blood was collected on the longitudinal cohort as mentioned below (**Figure 1**). Tissues were harvested with IACUC approved humane euthanasia at 90 dpi, 160 dpi, and endpoint (approximately 230 dpi). The experimental design is detailed in **Figure 1**. The early timepoint was classified at 90 dpi, midpoints spanned from 120 to 190 dpi, and endpoints were assigned to the 200 to 230 dpi timeframe.

**Figure 1:**
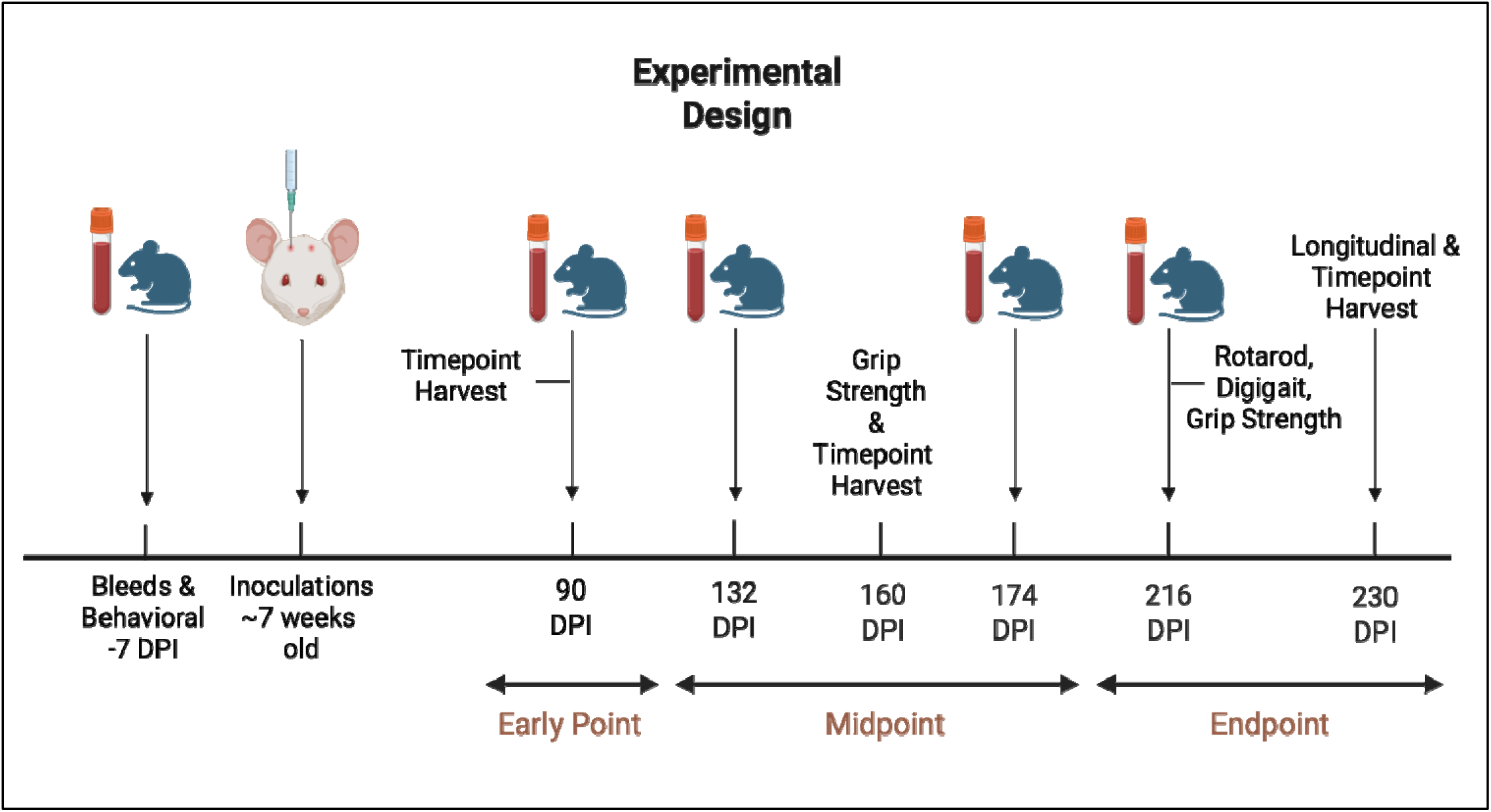
Experimental design of the animal portion of study. Various neurobehavioral assessments, blood collections, and harvests are shown at the appropriate timepoints. Early timepoints are classified at 90 dpi, midpoints span from 120 to 190 dpi, and endpoints are 200 to 230 dpi. Observational behavioral assessments occurred at the same timepoints as blood collections, other behavioral assessments are specified. Mice were harvested at disease endpoint, ∼230 dpi.

### 2.2 Submandibular Blood Collection and Processing

Less than 10% of total blood volume (based on mouse weight) was collected every 6 weeks via the submandibular blood collection method. A 5mm lancet (VWR, MSPP-GR5MM) was used to incise the submandibular (facial) vein as previously described^24^ and stored in K2 EDTA anticoagulant blood tubes on ice for less than 15 minutes. The samples were centrifuged at 14,000 rotations per minute (rpm) and plasma removed from the whole blood and stored at -80°C for future use in experiments.

### 2.3 Brain Homogenate Preparation

The brain homogenates used in the study were prepped using endpoint hippocampal and cortex tissues that were microdissected from the left hemisphere upon harvest^25–27^. The samples were weighed and diluted to 10% w/v using in-house RIPA buffer (25mM TRIS, 150mM NaCl, 0.1% (w/w) SDS, 0.5% (w/w) sodium deoxycholate, 1% (w/w) Triton X-100) with a protease inhibitor cocktail (Sigma Aldrich, #11836170001). Tissues were homogenized in 5E Pro bullet blender (Next Advance) on speed 20 for 60 seconds, aliquoted, and stored at 80°C.

### 2.4 Observational Neurobehavioral Profiling Assessments

Observational neurobehavioral profiling assessments were completed using a technique based on a modified SHIRPA evaluation^28–30^ at multiple time points (90,132,174, and 216 dpi). The scoring system was based on a four-point scale of sign severity; 0=no sign, 1=barely noticeable, 2=moderate, 3=severe with details of each observational behavioral scoring assessment provided in **Table S1**. The behavioral assessments were blindly conducted to maintain consistency and reduce bias.

### 2.5 Grip Strength

Grip strength testing was performed using a grip strength meter (Maze Engineers; #DS2-50N) on Tg1536 mice at 160 dpi and 200 dpi. Mice were placed on the meter’s grid in a vertical position until all four paws gripped the grid. The force was measured in (gF)/seconds while mice were gently guided downward steadily, allowing gravity to assist, until all paws released. This was repeated in two trials with a rest in between per mouse. Data was normalized to the mouse’s body weight in grams (g).

### 2.6 Rotarod

Motor coordination was assessed using a rotarod apparatus (Maze Engineers) on Tg1536 at 216 dpi. Mice were placed facing forward on the rotating bar, with the program set to run for 5 minutes accelerating from 4 to 40 rpm at a rate of 10r/m^2^. Testing occurred over the course of three days with three trials per day allowing the mice at least a 5-minute break between each trial. The latency to drop was automatically recorded by sensors in the machine. Mice housed in the same cage were tested at the same time. Additional guidelines for this test were considered as referenced^31^.

### 2.7 DigiGait^TM^

A detailed gait analysis was performed on the Tg1536 CWD and NBH longitudinal cohorts at 90dpi, 153dpi, 200dpi, and 230dpi using the DigiGait^TM^ treadmill manufactured by Mouse Specifics Inc. The DigiGait^TM^ incorporates Ventral Plane Imaging (VPI) to record mouse paw prints while they run at a fixed speed on a transparent treadmill belt. Each mouse was acclimated to the DigiGait^TM^ chamber for at least 30 seconds. After acclimation, the treadmill was sped up to 30cm/s and maintained for three to six seconds of consistent running. Although methods are similar to other references, this speed was altered from previous references^32–35^ after troubleshooting to better accommodate a comfortable speed for the Tg1536 mouse model. If mice were not able to run for three seconds, the longest obtained video was saved. After a successful run, mice were returned to their home cage. The analysis included 42 parameters reported by the DigiGait^TM^, although only 16 were included in results section. These were selected by including the parameters in which one or more were significantly different between CWD and NBH groups of mice at any timepoint or limb. The 26 parameters that were not significantly different between the two groups of mice were not reported, such as limb shared stance time, gait symmetry, overlap distance, among others. P-values were calculated using the student’s t-Test in Microsoft excel (version 16.110).

### 2.8 Histology

All tissues included in the study were processed as follows unless specifically stated. Mouse brain hemispheres were harvested at timepoints 90 dpi, 160 dpi, and 230 dpi and fixed in 10% formalin (Azer Scientific, PFNBF-20) for at least 48 hours^25,26,36^. For prion decontamination purposes, tissues used for histology were treated as recommended by the Biosafety in Microbiological and Biomedical Laboratories (BMBL) 6^th^ edition reference^36^. The mouse brain hemi-sections were immersed in 95% formic acid (Sigma-Aldrich, 1002640100) for 1 hour with gentle agitation in a biosafety cabinet, then decanted and replaced with PBS (Millipore, 6505-4L) + 0.02% Sodium Azide (Sigma-Aldrich, S8032). All tissues were then prepared under standard histologic tissue dehydration and paraffin embedding conditions^37^. Sagittal sections were taken at 5 μm, mounted on positively charged slides (Avantor, 48311-703), and stained with hematoxylin (VWR, 95057-844) and eosin (VWR, 95057-848) (H&E) to assess the morphology of the tissues using standard staining techniques^36^.

### 2.9 Plasma Biomarker Analysis

The Mesoscale Discovery System (MSD) MESO QuickPlex Q 60MM was used with the S-PLEX Neurology Panel 1 kit (#K15639S, Rockville, MD) to detect GFAP, NfL, and t-Tau in 2-fold diluted plasma at 90dpi, 132dpi, 174dpi, and 230dpi. Each sample was performed in duplicate, and product protocols were followed to improve accuracy and repeatability of results. Calibrations were performed according to product recommendations and standard curves that were utilized for each biomarker and timepoint are shown in (**Figure S1**). Quantitative data analysis was performed using Methodical Mind Enterprise Analysis Software (Meso Scale Discovery, Rockville, MD). Analyte concentrations for the MSD S-PLEX Neurology Panel 1 kit were interpolated from calibration standards using a 4-parameter logistic (4PL) regression curve-fitting model. The lower and upper limits of detection (LLOD and ULOD) for the standard curve were determined using a threshold of 2.5 standard deviations above the background (bottom of the range) and below the plateau (top of the range), respectively. Standard curves achieved a coefficient of determination (R^2^) of 0.999. Individual plasma concentration datapoints were calculated by averaging the two replicates. All datapoints were reported as the calculated concentration mean of two replicates and were within detection and fit curve range with a few exceptions described in the results.

### 2.10 RT-QuIC

RT-QuIC was performed using the QuIC kit from Priogen (Priogen Corp, St. Paul, MN) that used truncated prion protein AA-90-231 Syrian hamster substrate (0.1mg/ml, MNPROtein™). The assay was performed in accordance with product protocol. Brain homogenates (10%) were diluted 100-fold using the sample dilution buffer provided by Priogen. Reactions were performed in quadruplicate in a 96-well plate (Nunc, Thermo Fisher) with each well containing 2 µl of sample and 98 µl of RT-QuIC assay mix. The assay was incubated in a FLUOstar omega plate reader (BMG Labtech) at 42°C for 36 hours with double orbital shaking cycles of 1 min on and off at 700 rpm. The fluorescence was measured every 40 minutes with a gain of 1600, excitation of 448nm and emission of 482nm^38^. We established a cutoff for RT-QuIC analysis at 24 hours because after that timepoint, negative controls began to show ThT signal. RT-QuIC data after that timepoint was not included in the results section. The max point ratio (MPR) was calculated for each sample by dividing the maximum RFU value within 24 hours by the background RFU (average RFU of cycles 2-5). Samples with 3-4 replicates above the MPR threshold were considered RT-QuIC positive^39,40^.

### 2.11 Statistical Analysis

Graph Pad Prism 10 for MacOS version 10.6.1 and Microsoft® excel for Mac version 16.105.2 was used to perform the statistical analyses for all experiments.

## 3. Results

### 3.1 Observational Neurobehavioral Profiling Assessment

The observational neurobehavioral profiling assessments were grouped by four general behavioral categories. These were: 1) overall condition and appearance; 2) response to environment; 3) neurological signs and 4) motor ability. For detailed descriptions of the assessments under each category and scoring parameters see **Table S1**. Within the overall condition and appearance category, body weight was assessed over time by sex. As suggested by the name, CWD is known to cause significant weight loss, a sign that was exhibited in the diseased group in both sexes of mice **(Figure 2A&B).** The females had significantly heavier body weights than the NBH controls at both early and mid-timepoints, however, they were not significantly different at endpoint (**Figure 2A**). This is due to the notable weight loss in the female CWD mice at endpoint, who collectively lost 17% of their body weight from 216 to 230 DPI **(Figure 2A).** The males were not significantly different between the CWD and NBH cohorts until right before endpoint, however, average body weight began to decrease at midpoint 174 dpi (**Figure 2B**). In the response to environment category, nesting was impaired in the CWD mice, but this occurred at endpoint (**Figure 2C**). The neurological signs category revealed the Deer PrP-225SS CWD mice exhibited an observable phenotype in early midpoint, the tail hyperextension (**Figure 2D)**. This rigid tail elevation phenotype is significantly different between CWD mice and the NBH controls beginning as early as 132 dpi, prior to other observable behavioral signs of disease emerge, see **Figure 2D**. Trunk curl (within the motor ability category) and clasping (neurological signs category) were significantly different between the CWD and NBH cohorts at endpoint (**Figures 2E&F**).

**Figure 2:**
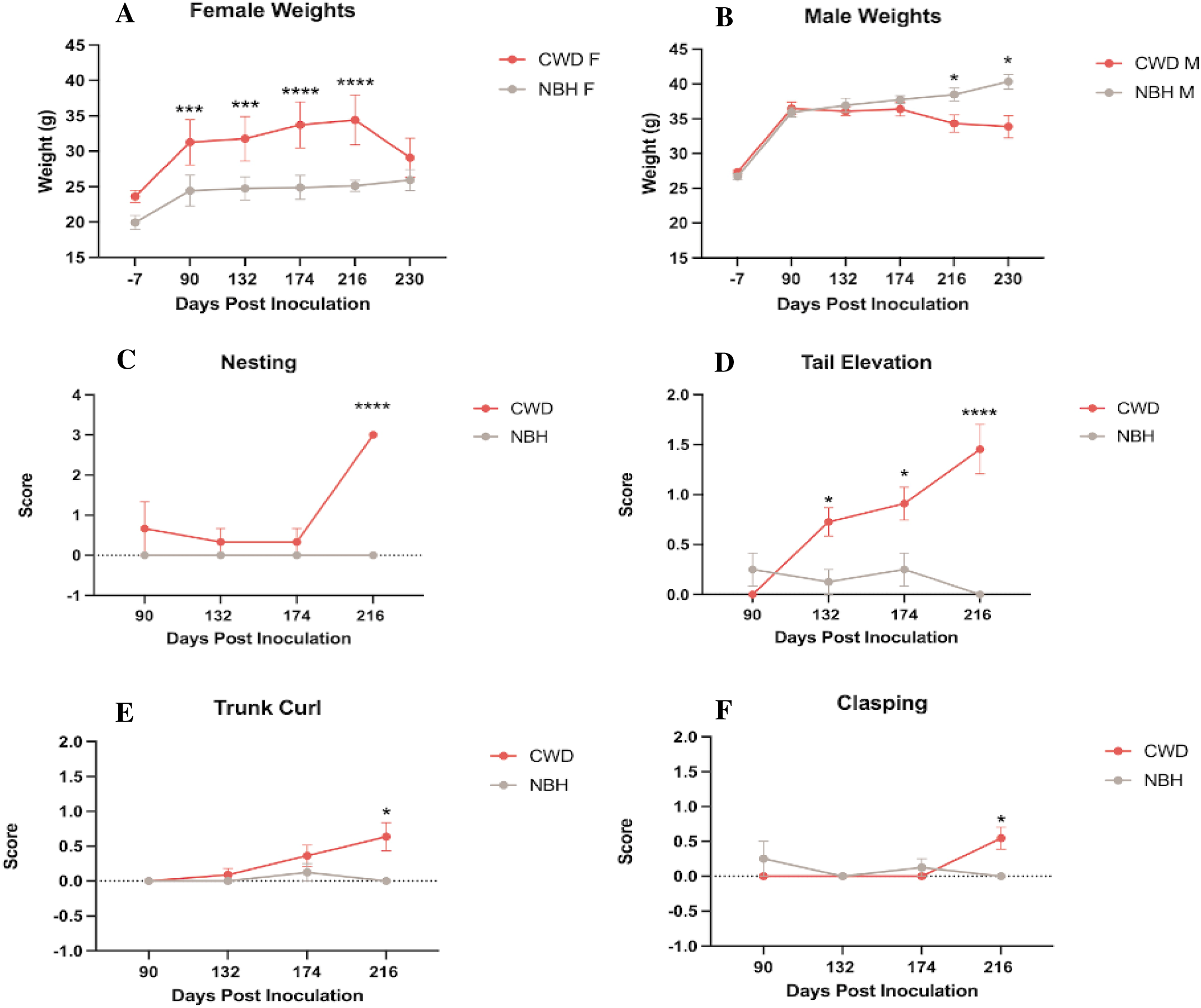
Representative observational neurobehavioral profiling in CWD Tg1536 mice displaying significantly different phenotypes between CWD mice and NBH controls over time. A) Average female weight (mean in grams (g)); B) Average male weight (mean in grams); C) Average nesting score; D) Average tail elevation score; E) Average trunk curl score; F) Average clasping score. CWD N=11 (5F, 6M) NBH N=8 (4F, 4M). Statistics performed as 2-Way ANOVA with Šídák’s multiple comparison test. P-values indicate *=<0.05, **<0.01, ***<0.001, ****<0.0001.

Remarkably, 30 of the observable neurobehavioral assessments performed were not significantly different between the groups at any timepoint including ataxia, coat condition, tremor, myoclonus, lordokyphosis, and hind limb paresis; some of which have been reported in other models of prion diseases such as scrapie^41,42^. A complete list of observable traits that were not found to be significantly different between the CWD and NBH mice can be viewed in **Table S2**.

### 3.2 Grip Strength Assessment

The Tg1536 mice were tested for grip strength at a midpoint (∼160 dpi) and endpoint (∼200 dpi). At both time points, overall differences can be observed in patterns of force and time between the CWD and NBH cohorts of mice. The two sexes also exhibited distinct patterns within the CWD group.

The beginning average force of the CWD females was 2.5gf/g, almost 3 times higher than the force of the NBH females (1gf/g). The two groups exhibited force for a similar amount of time, at 48 seconds and 46 seconds respectively (N=9, CWD=5, NBH=4) (**Figure 3A**). At endpoint, both CWD and NBH female groups began with a force of 2.2gf/g and exhibited force for a similar amount of time, at 42 seconds (**Figure 3B**). The average grip strength force patterns differed in the females when comparing the two different timepoints. At midpoint, the CWD female grip strength patterns visually appear distinct and higher than the NBH pattern, however, by endpoint, the two groups had similar patterns. Notably, the mean time of exhibiting force was longer in the midpoint at 47 seconds, then in the endpoint at 42 seconds (**Figure 3A&B**).

**Figure 3.**
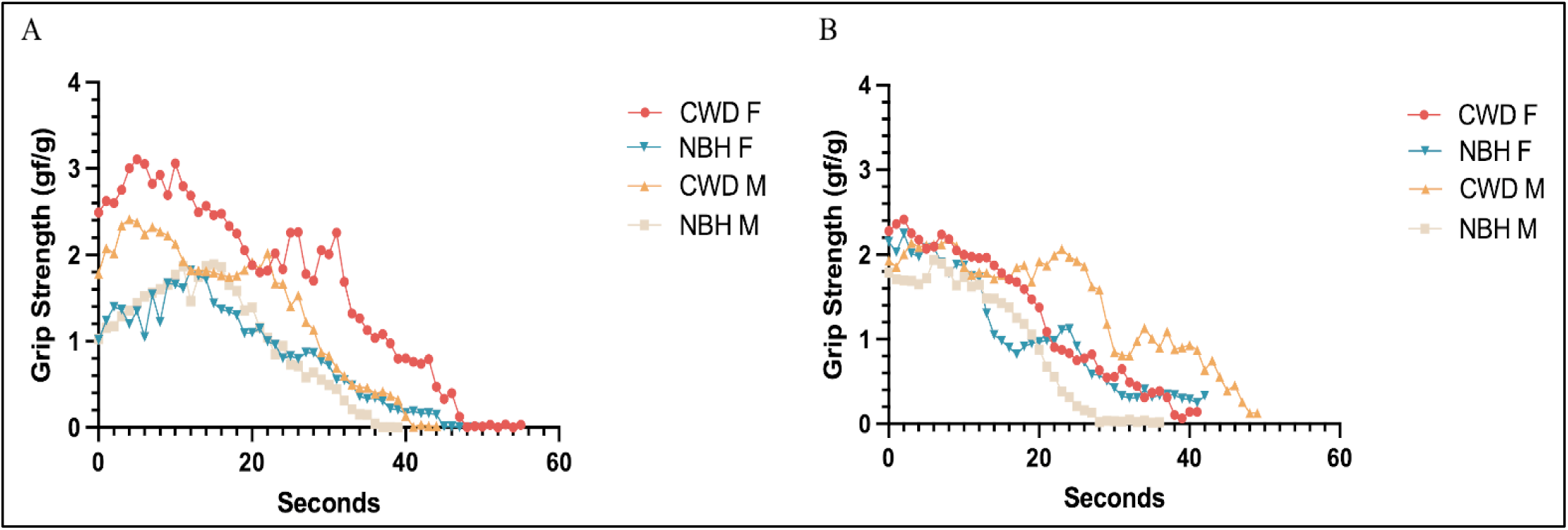
Average grip strength patterns exhibiting that CWD mice have higher and longer grip force than NBH controls beginning at disease midpoint (160 dpi) and remain at endpoint (200 dpi). [A] Grip strength patterns of force and time in both sexes of CWD and NBH mice at midpoint (160 dpi). [B] Grip strength patterns of force and time in both sexes of CWD and NBH mice at endpoint (200 dpi). CWD females, N=5 (red circles); NBH females, N=4 (blue upside-down triangle); CWD males, N=4 (orange triangles); NBH males, N=4 (tan squares).

CWD male mice exhibited a beginning average force of 1.8gf/g, almost double than the beginning force of the NBH males at 1gf/g. The CWD mice had a longer grip force time than the NBH controls at 42 seconds and 36 seconds, respectively (N=8, CWD=4, NBH=4) (**Figure 3A**). When comparing CWD to NBH males at endpoint, the mice began at the same force of 1.8gf/g, similar to that of the females, yet time spent exhibiting force was longer in the CWD males (48 seconds) than the NBH controls (28 seconds) (**Figure 3B**). Unlike the females, the average grip strength force patterns remained similar in the males when comparing the two different timepoints with the CWD males exhibiting visually higher and longer force patterns than the NBH group in general. Notably, at midpoint, both CWD females and males begin at a force being almost triple and double of their respective NBH counterparts, while all ending within seconds of each other. By endpoint, however, the pattern shifts; in all groups having similar starting forces but the males having more distinct and separate trial times.

In addition to the grip strength patterns, we analyzed the Tg1536 CWD and NBH mean grip strength force. The CWD group exhibited a higher average force than the NBH, consistent with the grip strength pattern results. At midpoint, the CWD group had a significantly higher mean force in 1/3 trials (Trial 1, p=0.0148) with the other trials trending higher in the CWD animals (**Figure 4A**). By endpoint, the CWD mice had a significantly higher mean force in 2/3 trials (Trial one, p=0.0099 and Trial three, p=0.0142) (**Figure 4B**) (CWD=9, NBH=8 in both midpoint and endpoint trials).

**Figure 4.**
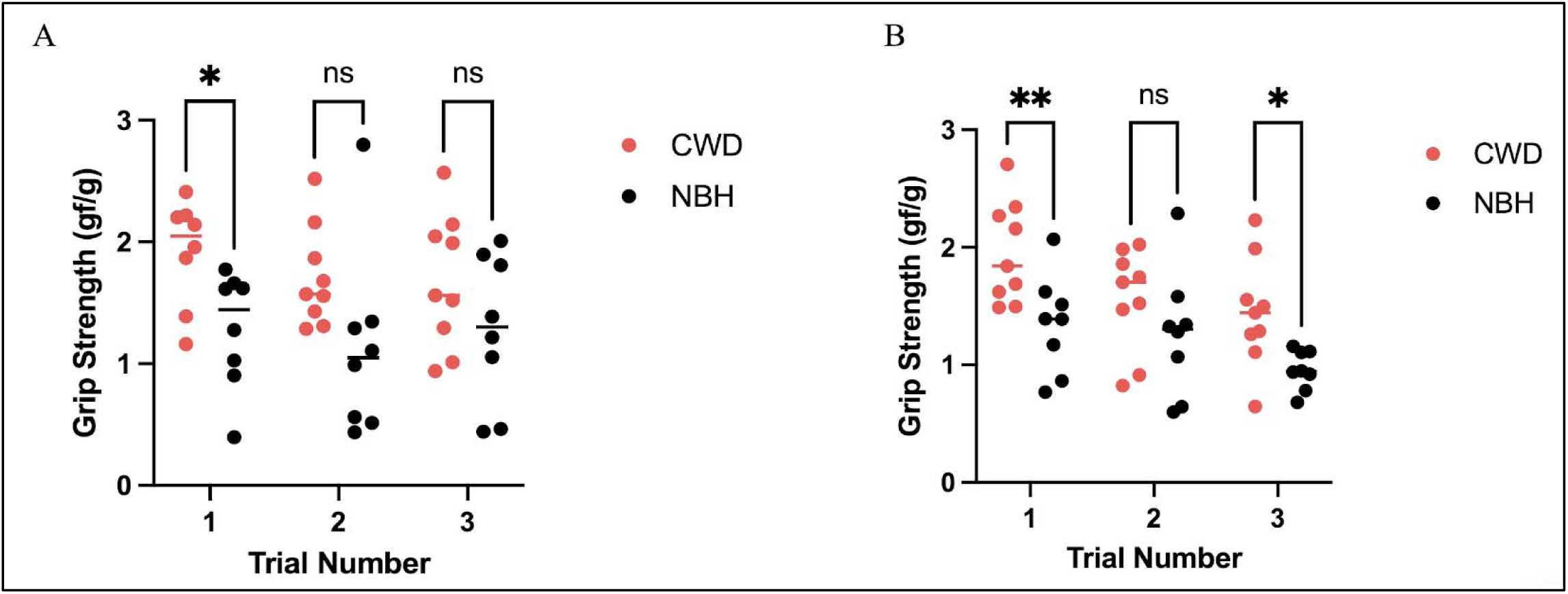
CWD mice begin to display significantly higher grip strength force than NBH controls at midpoint and remain higher at endpoint. **[A]** At midpoint (160 dpi), the mean force by individual CWD mice (red) are statistically higher than the NBH controls (gray) in trial 1 (p=0.0148) and are trending higher in trials 2 and 3. **[B]** By endpoint, the CWD mice have a statistically higher mean force than NBH in trial 1 (p=0.0099) and 3 (p=0.0142) with trial 2 trending higher. Statistics performed as 2-way ANOVA with Tukey’s multiple comparisons. (N=17, CWD=9, NBH=8).

### 3.3 Rotarod Assessment

In addition to the neurobehavioral and strength abnormalities previously discussed, coordination impairments appear in CWD mice as demonstrated in the rotarod assay at endpoint (∼216 dpi) (**Figure 5)**. The CWD mice displayed a shorter median latency, i.e., fell off of the rotating rod sooner, than the NBH mice in all nine runs with a significant difference on day 4 (p=0.0337), 6 (p=0.0264), 7 (p=0.0162), and 8 (p=0.0168) **(Figure 5A)**.

**Figure 5:**
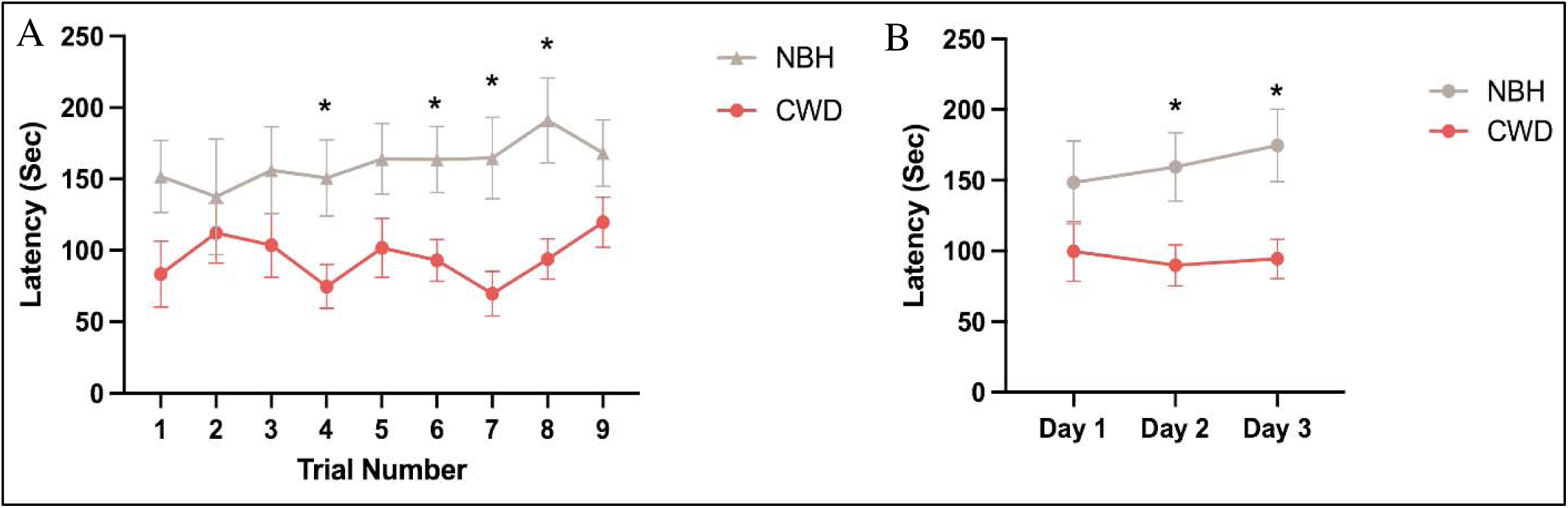
CWD mice display coordination and skill acquisition deficits as opposed to NBH controls in the rotatod assay at endpoint (∼216 dpi). **[A]** Latency median time (±SEM) during each trial (trial number 1-3=day 1, 4-6=day 2, 7-9=day 3). [B] Latency median time (±SEM) by day. Red represents CWD mice (n=9) and gray represents NBH mice (n=7). Significance (p < 0.05) denoted in asterisks and was calculated using 2-Way ANOVA with Sidak’s multiple comparison test.

Furthermore, NBH mice had a slight increase in their median latency over the course of the three days while the CWD mice had a slight decrease. Because normal mice are expected to improve on the rotarod over the three-day trial, these findings suggest that CWD mice have deficits in skill acquisition in addition to impaired motor coordination **(Figure 5B).**

### 3.4 DigiGait^TM^ Assessment

DigiGait^TM^ data revealed that hind limb phenotypes were present in early and mid-timepoints, while the front limb parameter differences were more often seen in late timepoints **(Table 2)**. A higher number of gait phenotypes (11/32) were significantly different between CWD and NBH mice when combining early and mid-timepoints (90 and 153 dpi) in the hindlimb, while only 7/32 were significantly different in the front limb at those timepoints **(Table 2)**. In late timepoints (200 and 230 dpi), the hindlimb had only 4/32 significant parameters, while the front limb had 10/32 parameters that were significantly different between the CWD and NBH mice **(Table 2)**. Notably, the only markers that were present at early or mid-timepoints and remained until late timepoints were hindlimb brake, front limb % swing stride, front limb stance/swing, and both hind and front limb paw angle variability. We hypothesized ataxia coefficient would be significantly elevated in the CWD mice as compared to controls and progressively increase as disease progressed, however, the results revealed it was only significantly different at earlier timepoints in both limbs.

**Table 2:**
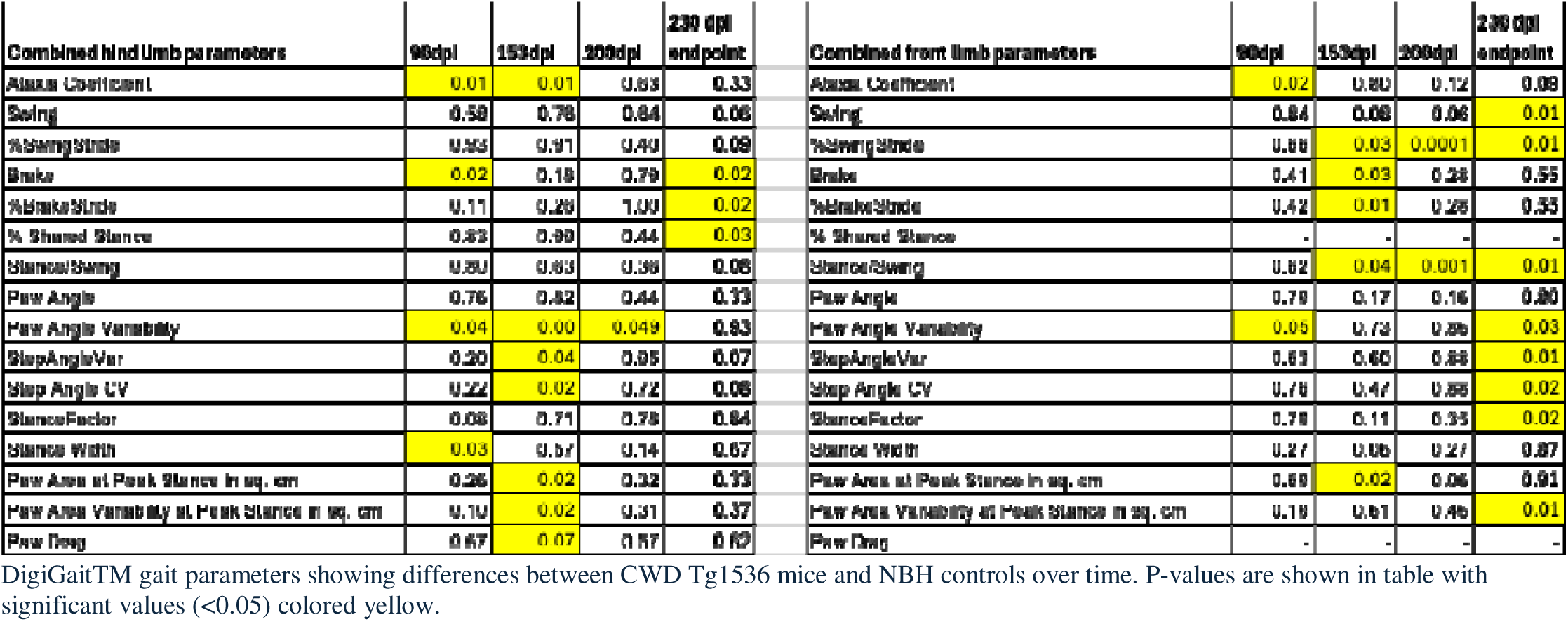
CWD mice display subtle gait phenotypes in both hind and front limb at all timepoints.

### 3.5 Plasma Biomarkers of CWD

Previously identified to be effective plasma biomarkers in other neurodegenerative diseases^43–51^, GFAP, NfL and t-Tau were analyzed in the plasma samples of Tg1536 CWD and NBH inoculated mice. All three biomarkers progressively increased mean concentration in the CWD plasma as opposed to NBH controls over time (**Figure 6**).

**Figure 6:**
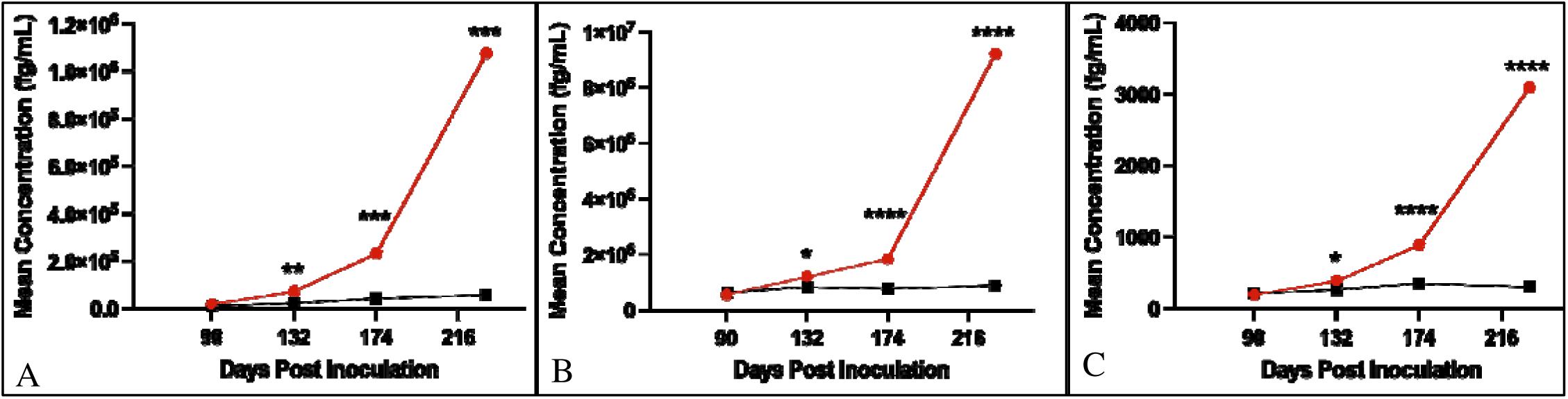
GFAP, NfL, and t-Tau mean plasma concentrations elevate over time in CWD Tg1536 mice. **[A]** Mean concentration of plasma GFAP displayed significant elevation in CWD mice (red) compared to NBH (gray) beginning at early midpoint (132 dpi) (p=0.0014); **[B]** Mean concentration of plasma NfL displayed significant elevation in CWD mice beginning at midpoint (174 dpi) (p=<0.0001); **[C]** Mean concentration of plasma t-Tau displayed significant elevation in CWD mice beginning at midpoint (174 dpi) (p=<0.0001). P-values indicate *=<0.05, **<0.01, ***<0.001, ****<0.0001. Statistics performed as 2-way ANOVA with Tukey’s multiple comparison (n=20, CWD=12, NBH=8).

When individual plasma samples were analyzed by time, our early timepoint of 90 dpi did not display statistical differences in the median concentration between the CWD and NBH groups in any of the three biomarkers, however, the medians were already trending slightly higher in the CWD group (**Figure 7**). The GFAP concentration range (fg/ml) was 4,308 to 40,192 with a median of 8,626 (n=9) in the Tg1536 NBH plasma, while the CWD plasma had a concentration range (fg/ml) of 9,805 to 35,823 with a median of 16,650 (n=14) (**Figure 7A**). The NfL concentration range was 204,685 to 1,631,950 with a median of 477,320 in the NBH (**Figure 7B**), while CWD had a range of 372,676 to 672,340 with a median of 527,123 (**Figure 7B**). Lastly, t-Tau had a concentration range of 60 to 548 with a median of 170 in NBH, and a range of 144 to 279 with a median of 190 in CWD (**Figure 7C**).

**Figure 7:**
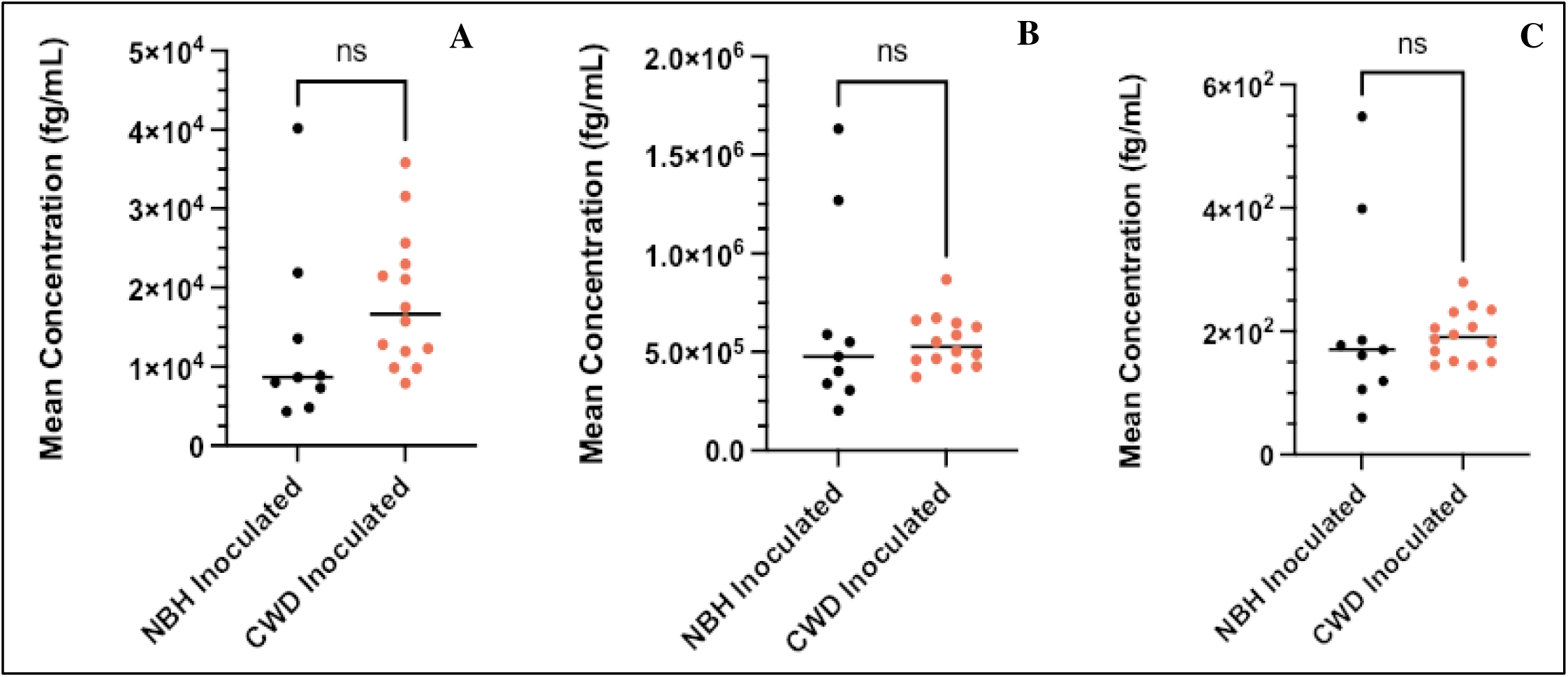
GFAP, NfL, and t-Tau plasma concentrations are not significantly different at the early timepoint (90 dpi). **[A]** Median GFAP plasma concentration appears to be higher in CWD, although not statistically significant; **[B]** Median NfL plasma concentration appears to be slightly higher in CWD, although not statistically significant; **[C]** Mean t-Tau plasma concentration appears to be slightly higher in CWD, although not statistically significant. Individual datapoints indicate the mean of two replicates per sample. Median is indicated by the horizontal bar. CWD shown in red and NBH shown in black. Statistics performed as Welch’s T-test (n=23, CWD=14, NBH=9).

By 132 days post inoculation, statistically higher levels of all biomarkers can be seen in CWD plasma (**Figure 8**). For GFAP, concentrations in the NBH group ranged from 7,610 to 46,637 fg/ml with a median of 22,728 (n=10) (**Figure 8A**). In contrast, the CWD group exhibited significantly higher GFAP concentrations, ranging from 22,271 to 240,460 fg/ml with a median of 55,377 (n=14; p=0.0064) (**Figure 8A**). For NfL, the NBH group had a concentration range of 374,945 to 1,583,284 fg/ml with a median of 813,650 (n=10) (**Figure 8B)**, while the CWD group had a higher range of 556,266 to 2,010,382 fg/ml with a median of 1,180,000 (n=14; p=0.0461) (**Figure 8B**). Although t-Tau was detected at lower absolute concentrations, the values were also higher in the CWD group. The NBH group ranged from 143 to 443 fg/ml with a median of 249 (n=10), whereas the CWD group ranged from 192 to 607 fg/ml with a median of 384 (n=14; p=0.0215) (**Figure 8C**).

**Figure 8:**
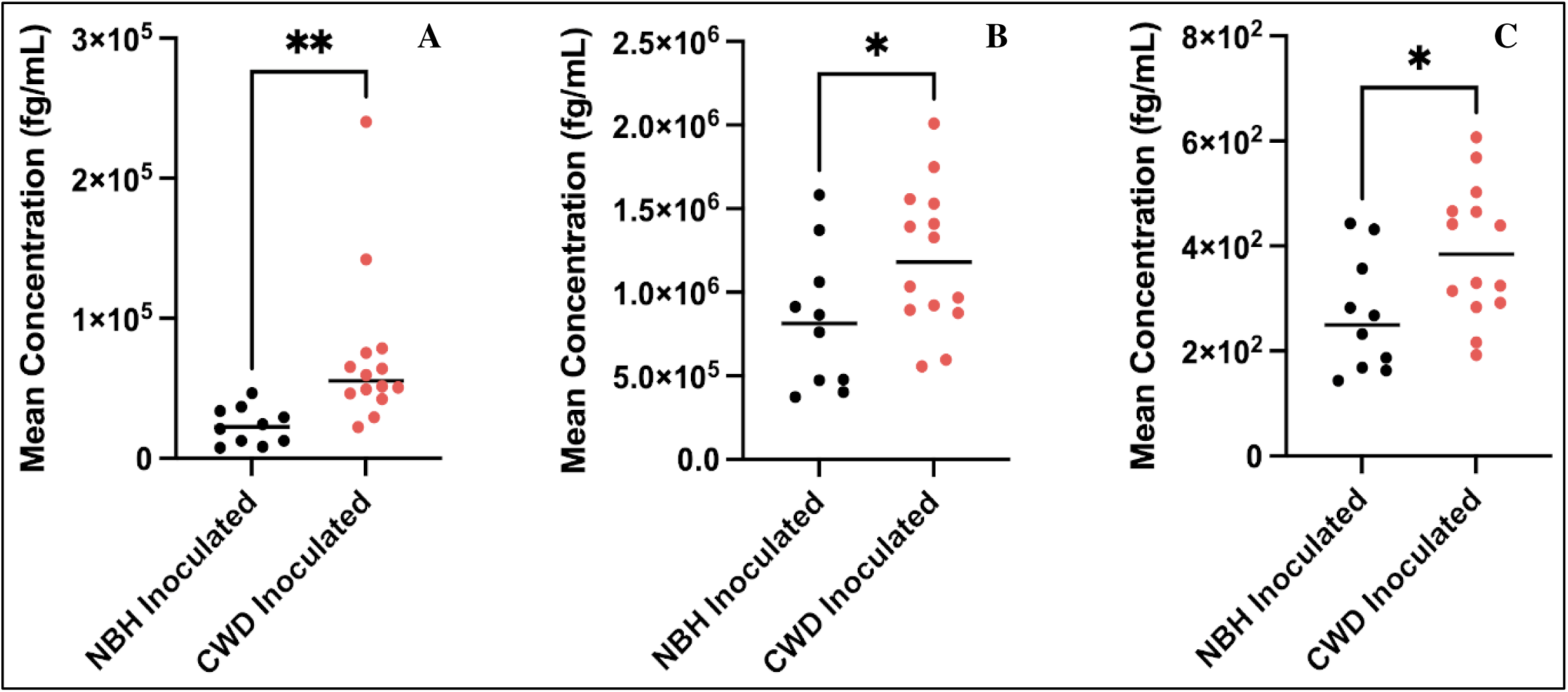
GFAP, NfL, and t-Tau plasma concentrations are beginning to be significantly elevated at an early midpoint (132 dpi). **[A]** Median GFAP showing significantly higher concentrations in CWD plasma (p=0.0064); **[B]** Median NfL plasma concentration showing significantly higher concentrations in CWD plasma (p=0.0461); **[C]** Mean t-Tau plasma concentration showing significantly higher concentrations in CWD plasma (p=0.0215). Individual datapoints indicate the mean of two replicates per sample. Median is indicated by the horizontal bar. CWD shown in red and NBH shown in black. Statistics performed as Welch’s T-test (n=24, CWD=14, NBH=10).

By the next mid timepoint (174 dpi), all three biomarkers showed strong statistical elevation in the CWD plasma concentration (**Figure 9**). The NBH plasma had a GFAP concentration (fg/ml) range of 22,374 to 94,138 with a median of 34,075 (n=8) (**Figure 9A**), whereas the CWD group had a significantly higher mean concentration (fg/ml) range of 90,815 to 499,520 with a median of 182,180 (n=12; p=0.0005) (**Figure 9A**). The NBH plasma had an NfL concentration range of 621,480 to 937,892 with a median of 744,932 (n=8) (**Figure 9B**), while CWD had a significantly higher range of 1,009,350 to 2,876,696 with a median of 1,757,000 (n=12; p=<0.0001) (**Figure 9B**). The t-Tau biomarker at this timepoint had an incredibly strong significant elevation in the CWD plasma (p=<0.0001). The NBH group had a mean concentration range of 280 to 518 with a median of 326 (n=8) (**Figure 9C**), and the CWD group had a mean concentration range of 455 to 1519, resulting in a median that was over double that of the NBH at 871 (n=12) (**Figure 9C**).

**Figure 9:**
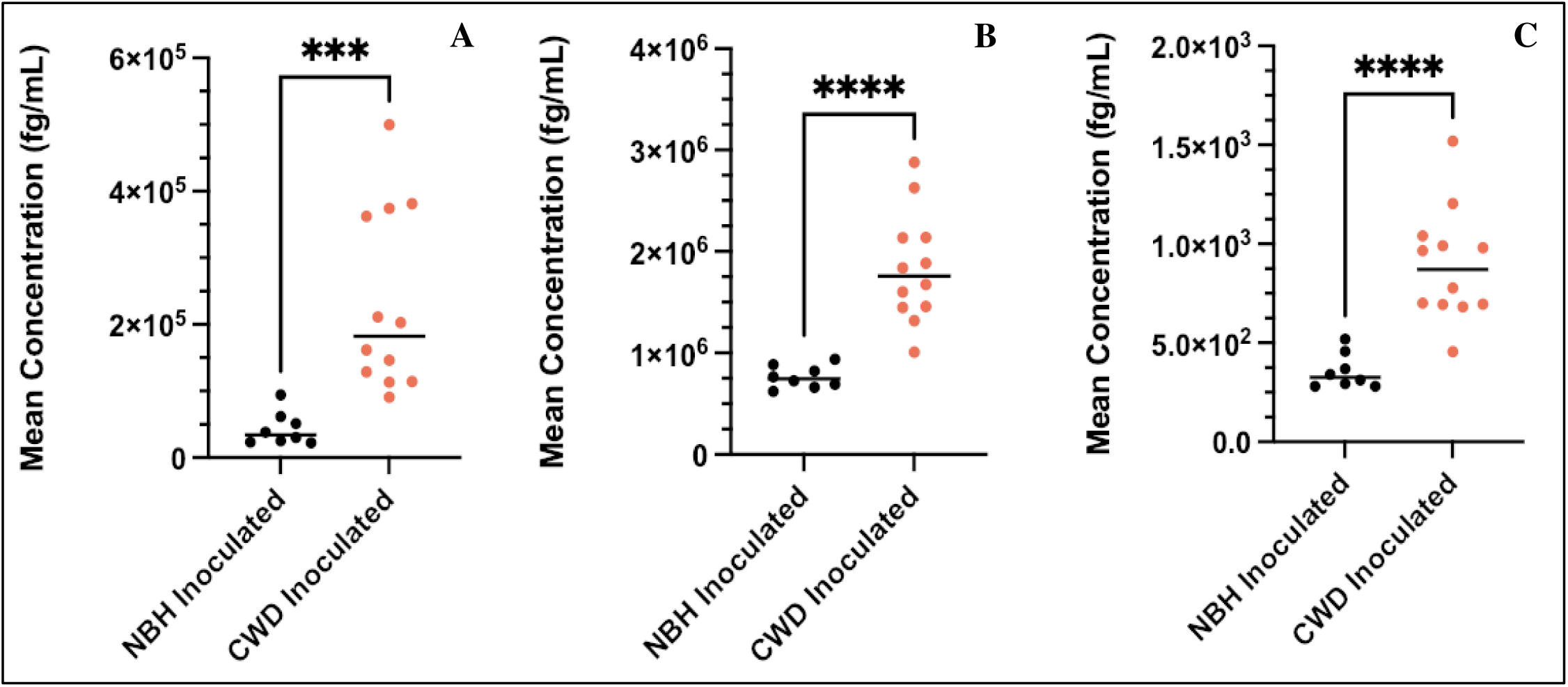
GFAP, NfL, and t-Tau plasma concentrations are significantly elevated at midpoint of 174 dpi. **A]** GFAP concentration significantly elevated in CWD plasma (p==0.0005); **[B]** NfL concentration significantly elevated in CWD plasma (p=<0.0001); **[C]** t-Tau concentration significantly elevated in CWD plasma (p=<0.0001). Individual datapoints indicate the mean of two replicates per sample. Median is indicated by the horizontal bar. CWD shown in red and NBH shown in black. Statistics performed as Welch’s T-test (n=20, CWD=12, NBH=8).

High concentrations of all three biomarkers were seen at endpoint (230 dpi) in the CWD plasma samples **(Figure 10)**. The GFAP NBH group had a concentration range of 13,138 to 88,205 with a median of 67,982 (n=8) **(Figure 10A),** and the CWD group had a mean concentration range of 317,467 to 2,441,973 with a median of 955,056 (n=14) (p=0.0002) **(Figure 10A).** The NBH plasma NfL ranged from 403,430 to 1,373,630 with a median of 834,905 (n=8), while the CWD group ranged from 3,102,739 to 12,938,767 with a median of 10,663,007 (n=14) (p=<0.0001) **(Figure 10B).** The NBH plasma t-Tau had a mean concentration range of 131 to 509 with a median of 271.5 (n=8), while the CWD group had a mean range of 973 to 4314 with a median of 3558.8 (n=14) (p=<0.0001) **(Figure 10C).**

**Figure 10:**
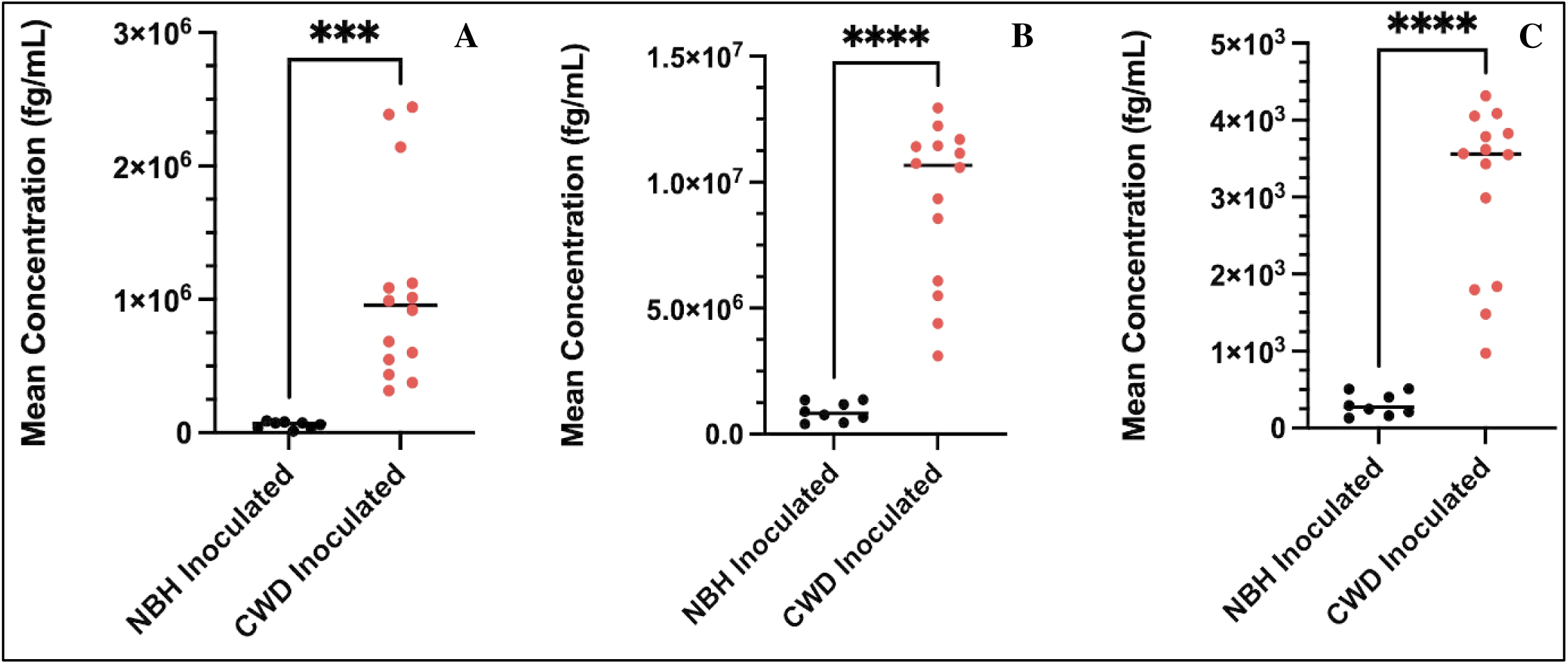
GFAP, NfL, and t-Tau plasma concentrations are significantly elevated at the late timepoint of 230 dpi. **[A]** GFAP concentration significantly elevated in CWD plasma (p==0.0002); **[B]** NfL concentration significantly elevated in CWD plasma (p=<0.0001); **[C]** t-Tau concentration significantly elevated in CWD plasma (p=<0.0001). Individual datapoints indicate the mean of two replicates per sample. Median is indicated by the horizontal bar. CWD shown in red and NBH shown in black. Statistics performed as Welch’s T-test (n=22, CWD=14, NBH=8).

The signal coefficient of variation (CV) was assessed for data quality, with the following scale classifications: under 10% reflected excellent repeatability and high precision; 10%-20% signified less precision yet is acceptable for most standard bioassays nearing the limit of detection; and above 25% indicated high variability and poor precision within the samples. All samples were the under 25% CV threshold, with averages in the excellent category, indicating that overall, our sample reproducibility was high. GFAP displayed an average CV of 7.83% (range of 2.07-17.20%), NfL had a 3.59% average (range of 0.043-8.55), and t-Tau exhibited an average of 6.12% (range of 1.43-12.46). All biomarker datapoints in the study were within detection and fit curve range with some exceptions in the endpoint group. In this group, 3 of 18 GFAP CWD positive samples reported above the fit curve range, whereas NfL had 10 of 18 CWD positive samples above the fit curve range. To maintain consistency in methods and dilution factors across all timepoints, these concentrations were retained and included in the statistical analyses. The CV in these particular samples demonstrated overall high reproducibility and precision of our samples, despite them being reported above the fit curve range. The three GFAP CV readings were 3.70, 5.01, and 14.4, while the ten NfL CV readings ranged from 0.04-5.84.

A set of controls were included on each plate consisting of the 5xFAD, uninoculated FVB/NJ, and uninoculated Tg1536 models. A representation of this data is included at endpoint **(Fig. S2).** Both replicates of each biomarker are displayed for individual mice to demonstrate concentrations in another neurodegeneration model (5xFAD), in uninoculated transgenic mice (Tg1536), and the background strain of the transgenic model (FVB/NJ). Represented on the same scale as endpoint NfL **(Fig. 10B),** most of the controls exhibited no concentration elevation, with the exception of NfL in the 5xFAD model which was expected. Notably, the close precision of each replicate is demonstrated in **Fig.S2**.

### 3.6 RT-QuIC

Conformation of CWD and prion seeding activity was performed in the Tg1536 CWD mice at endpoint (230 dpi). Sample numbers C1-C8 were Tg1536 CWD inoculated mice and N1-N8 were NBH at 230 dpi. Control 1 was *PRNP* null transgenic mouse brain, control 2 was the mule deer CWD positive brain homogenate used for inoculation of the study mice, and control 3 was a non-diseased normal mule deer brain homogenate used for inoculating the NBH control mice. Samples with an MPR above the threshold line at 2, indicated by the red dotted line, were considered positive. The Tg1536 CWD inoculated samples used in the study (C1-C8) were all confirmed to have a positive CWD status and prion seeding activity demonstrated by MPR (**Figure 11**). Despite samples C1 and C8 meeting overall positive status by MPR, not all replicates (3/4) were positive **(Figure 11 & Table 3)**. The lag phase was also longer on both sample C1 and C8 **(Table 3)**. These findings together suggest the seeding activity is lower than the other samples, however, still present. The seeding profiles of each sample via kinetic data is agreeable to MPR analysis as seen in **Figure S3A &B.**

**Figure 11:**
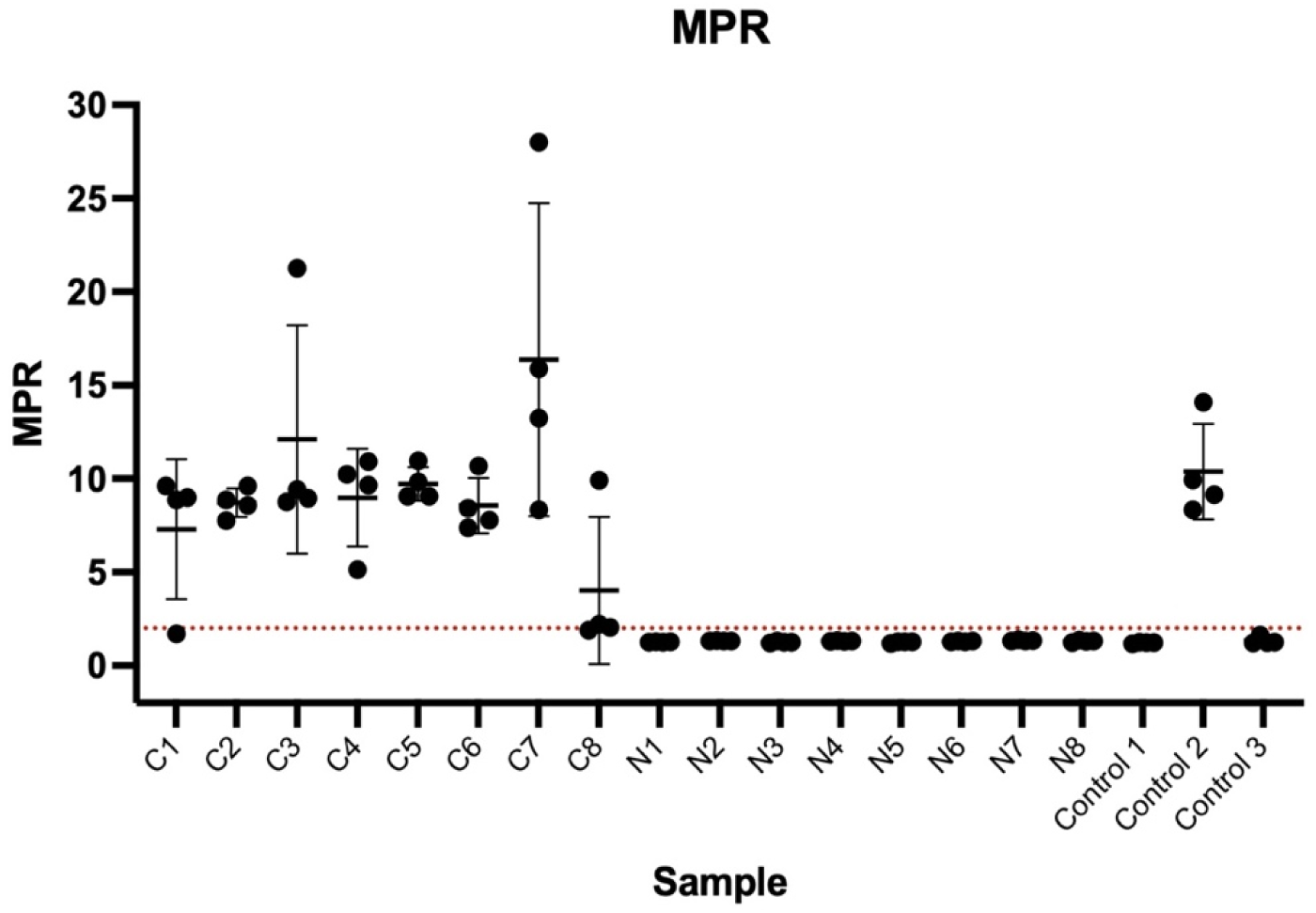
RT-QuIC assay demonstrating positive CWD status and prion seeding activity of inoculated Tg1536 mice at endpoint (230dpi) used in the study. Tg1536 CWD mice brain homogenates are above the MPR threshold (indicated by dotted red line) showing prion seeding activity, whereas the Tg1536 NBH brain homogenates show no prion seeding activity and are below the MPR threshold. Control 1-3 brain homogenates (from a *PRNP* null mouse, a CWD positive deer, a CWD negative deer respectively) were additional controls and are positive and negative as expected based on MPR. Each datapoint represents one RT-QuIC run and samples were performed in quadruplicates.

**Table 3:** RT-QuIC assay results validating prion seeding activity or lack of prion seeding activity in Tg1536 CWD and NBH inoculated mice used in the study.

| RT-QuIC Sample Number | Inoculation | Average MPR (Max Point Ratio) | Prion RT-QuIC outcome (ThT+/total wells (4)) | Lag Phase (h) |
| --- | --- | --- | --- | --- |
| C1 | CWD | 7.29 ± 3.7 | 3/4 | 18.6 ± 2.7 |
| C2 | CWD | 8.71 ± 0.8 | 4/4 | 13.2 ± 3 |
| C3 | CWD | 12.10 ± 6.1 | 4/4 | 11.8 ± 1.6 |
| C4 | CWD | 8.98 ± 2.6 | 4/4 | 17.7 ± 2.8 |
| C5 | CWD | 9.73 ± 0.9 | 4/4 | 17.7 ± 1.9 |
| C6 | CWD | 8.57 ± 1.5 | 4/4 | 13.7 ± 0.6 |
| C7 | CWD | 16.37 ± 8.4 | 4/4 | 7.1 ± 4.8 |
| C8 | CWD | 4.02 ± 3.9 | 3/4 | 21.2 ± 2.4 |
| N1 | NBH | 1.27 ± 0.0 | 0/4 | - |
| N2 | NBH | 1.32 ± 0.0 | 0/4 | - |
| N3 | NBH | 1.26 ± 0.0 | 0/4 | - |
| N4 | NBH | 1.31 ± 0.0 | 0/4 | - |
| N5 | NBH | 1.25 ± 0.0 | 0/4 | - |
| N6 | NBH | 1.30 ± 0.0 | 0/4 | - |
| N7 | NBH | 1.34 ± 0.0 | 0/4 | - |
| N8 | NBH | 1.30 ± 0.0 | 0/4 | - |
| Control 1 | N/A | 1.22 ± 0.0 | 0/4 | - |
| Control 2 | CWD | 10.38 ± 2.56 | 4/4 | 10.8 ± 0.1 |
| Control 3 | NBH | 1.33 ± 0.2 | 0/4 | - |

MPR and lag phase in Tg1536 CWD inoculated mice verifies CWD status and prion seeding activity along with absence of prion seeding activity in the NBH inoculated mice. RT-QuIC sample number C1-C8: Tg1536 CWD brain homogenates at 230dpi, N1-N8: Tg1536 NBH brain homogenates at 230 dpi, Control 1: PRNP null mouse brain homogenate, Control 2: CWD positive deer brain homogenate, Control 3: CWD negative deer brain. Assays were performed in quadruplicate.

All Tg1536 NBH samples were below MPR as expected, confirming the absence of prion seeding activity **(Figure 11 & Table 3)**. Control 1 served as a negative control and had the lowest MPR as expected; control 2 demonstrated prion seeding activity and validated our CWD inoculum used in the mice; control 3 lacked prion seeding activity and validated our NBH inoculum **(Figure 11 & Table 3)**.

### 3.7 Histology

We performed H&E staining and scored the pathologic spongiform degeneration present in early, middle, and late timepoints in disease to better understand the neurobehavioral and biomarker changes characterized previously in the study. Increasing degrees of spongiform degeneration were observed in the hippocampus and surrounding regions of CWD inoculated mice over the course of disease (**Figure 12**). The degree of spongiform degeneration was scored as minor in the CWD mice even at the earliest 90 dpi timepoint (**Figure 12A**). At the midpoint of disease progression (160 dpi), degeneration was scored as moderate, whereas by the endpoint (230 dpi), degeneration was severe compared with the NBH-inoculated Tg1536 control (**Figure 12B, C, &D)**.

**Figure 12:**
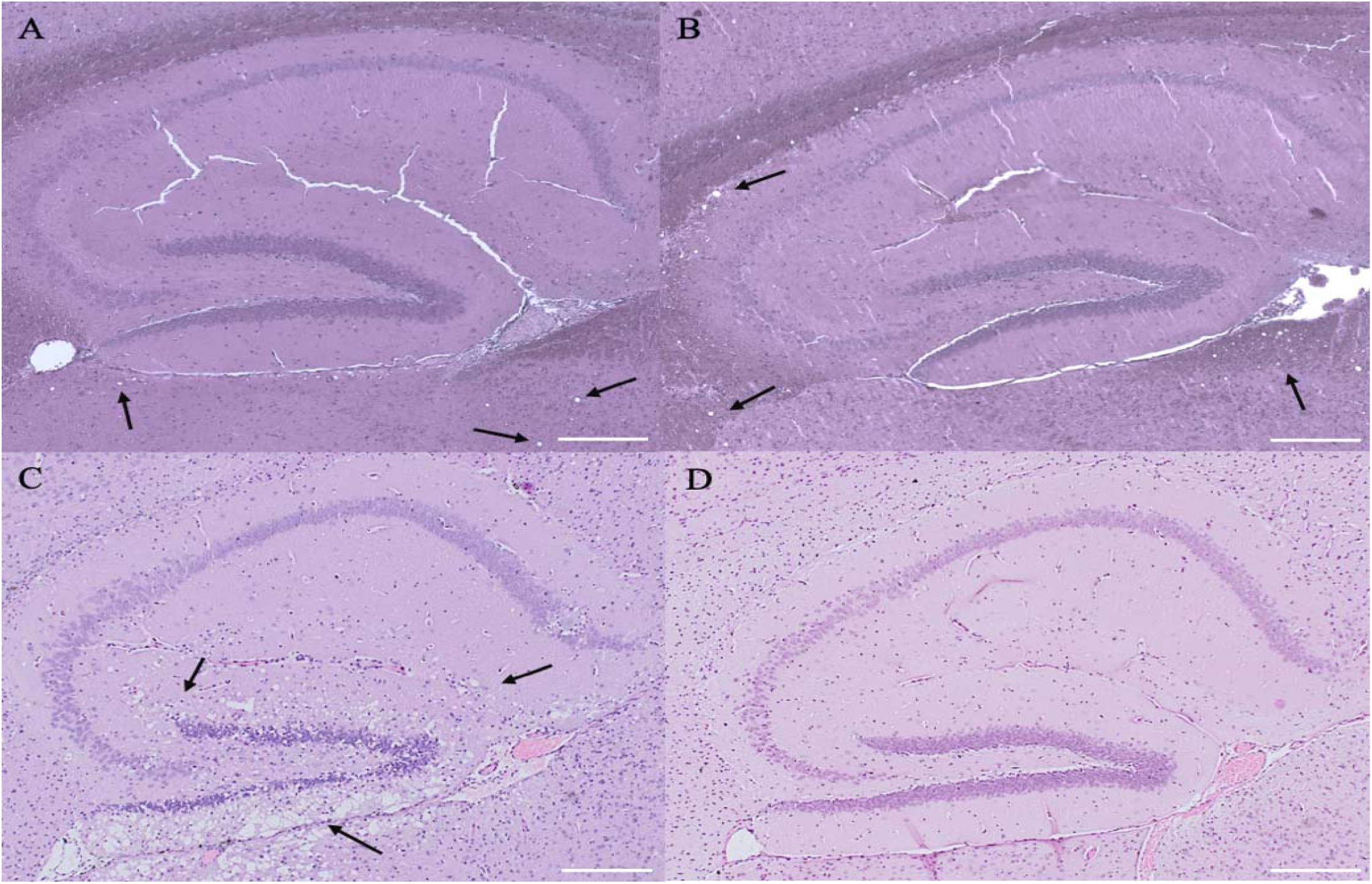
Increasing degrees of neuropathology occur over early, middle, and late CWD timepoints in the Tg1536 CWD mice. Representative examples of H&E staining of hippocampus in Tg1536 mouse brains over a time course. [A] Tg1536 CWD hippocampus at early timepoint 90dpi showing very little to no spongiform degeneration; [B] Tg1536 CWD hippocampus at midpoint 160dpi representing mild spongiform degeneration; [C] Tg1536 CWD hippocampus at endpoint (230dpi) with a higher degree of spongiform degeneration; **[D]** Tg1536 NBH hippocampus at endpoint (230dpi) lacking spongiform degeneration. Arrows indicate examples of spongiform degeneration. Scale bar = 200um

## 4. Discussion

Using phenotypic and biological markers, we report for the first time, the detection and elevation of NfL, GFAP, and t-Tau in plasma along with novel phenotypes in Tg1536 CWD mice over a complete disease time course. Biomarker levels, combined with phenotypic analysis allowed for full insight into the chronological progression of disease which uncovered signs of disease even at early antemortem timepoints.

### 4.1 Observational neurobehavioral profiling assessment

Historically, neurobehavioral assessments in mice have led to distinct advances in our understanding of other neurological diseases^16,17,37,52–55^. Our CPP methods shown to identify and characterize neuropathological timepoints may be useful in other applications such as to determine differing prion strains or evaluation of future therapeutic interventions. Importantly, the timing of therapy administration in prion diseases has been shown to be important to treatment efficacy^16^. Although studies document investigation of behavioral phenotypes of mouse models of other neurologic diseases^16–18^, there are few behavioral studies published regarding CWD in mouse models, and those are very limited^19,20^. Our CWD comprehensive phenotypic profiling evaluations have enabled the earliest and most complete assessment of behavior related disease markers, investigating a battery of over 40 characteristics relevant to neurodegenerative diseases in mouse models which identified unique phenotypes even at early timepoints. Remarkably, the behavioral assessments revealed the Tg1536 CWD mice exhibited spastic or rigid phenotypes such as tail hyperextensions opposed to the weakness scrapie models display^10,77^. This rigid tail elevation phenotype, which differs from plastic tail in scrapie mice^77^, has not been reported in previous literature of CWD mouse models, yet appears in the CWD mice at early midpoint, prior to all other signs of disease emergence, with the exception of the ataxia coefficients detected by the DigiGait^TM^. We predicted nesting behavior would emerge early in disease, as a previous study performed in-house demonstrated that nest building behaviors changed in scrapie infected mouse models as early as 90 dpi, around the same time that NfL levels began to increase^16^. Despite our incorrect prediction, our discovery of late nesting alterations is a meaningful finding as this is the first report of altered nesting behavior in CWD models. Similarly, trunk curling, cited in neurological diseases such as Huntington’s and Leigh syndrome^56,57^, has yet to be reported in CWD mouse models, however, these mice exhibited this sign as early as midpoint with significance at endpoint. While the phenotypic profiling revealed new and unexpected phenotypes and trends, it was unsurprising that the diseased mice showed weight loss at endpoint. Although both sexes lost weight by disease endpoint, the timing and patterns were different between sexes, as females lost weight at endpoint only, yet the males lost weight earlier in disease. CWD females had consistent weight gain over the course of disease, with the CWD cohort maintaining significantly heavier weights with higher gain than the NBH cohort. Weight loss began in CWD males at midpoint, however, prior to that were not significantly different between CWD and NBH groups.

### 4.2 Grip strength, rotarod, and DigiGait^TM^ assessments

In addition to the observational phenotypic profiling findings, we discovered unique phenotypes when grip strength, rotarod, and DigiGait^TM^ assessments were analyzed. While the rotarod test revealed an expected deficit in coordination in the CWD mice, the grip strength test results were unexpected. Tested at midpoint and endpoint, it appeared that the CWD mice exhibited higher grip strength patterns in both force and time in both sexes of mice. This was unexpected, as neurological dysfunction typically results in decreased grip strength^53,58–60^. These results, in addition to the clinical signs of tail rigidity and elevation, suggest a rigid phenotype might be occurring, or conversely, a loss of refinement when performing motor and strength functions. Brain regions and motor control pathways are known to be affected by spongiform change and neuron loss^1,61,62^, however, have not previously been linked with an increased grip strength. Future investigations need to be performed to further characterize the “rigid” phenotype and determine the exact etiology.

Interestingly, the highly detailed DigiGait^TM^ analysis revealed that the ataxia coefficient was significant only during early disease, despite ataxia being one of the few phenotypes consistently reported in CWD-affected wildlife, where it is typically observed later in disease progression^7,61,63^. This temporal discrepancy suggests that quantitative gait analysis may detect subtle early locomotor instability that does not persist as a distinct phenotype at later stages, potentially reflecting the emergence of more variable motor abnormalities as disease progresses. Out of 42 specific gait parameters measured on the DigiGait^TM^ for all four limbs, only a subset of 16 were significantly different between the two groups and often were measures of variability. The results were varied with few gait parameters having a distinct significance between groups as we predicted would occur based on our assumptions that gait perturbations would increase in CWD animals as disease progressed. Although these findings differed from our initial predictions, they are consistent with previous reports describing subtle and variable gait phenotypes, including ataxia, in CWD-affected wildlife^7,61^.

### 4.3 Plasma biomarkers of CWD

Practical testing modalities of CWD is crucial to reducing spread of disease. Currently, standard CWD detection methods rely on postmortem collection of difficult-to-obtain tissues such as brain stem or lymph nodes and require arduous laboratory-based techniques such as ELISAs, RT-QuIC, or IHC. Validation of less invasive sample types, such as blood, for early diagnostic testing represents an improvement over current tissue-based methods, including tonsil, rectal, or retropharyngeal lymph node biopsies, which may either lack sufficient sensitivity early in disease or require more invasive sampling^64^. Blood is considered an ideal sample type for many diagnostic assays because it is readily accessible; however, current blood-based testing capabilities for CWD remain limited, in part because the kinetics of prionemia across the disease course have not yet been fully characterized^64^. Although research is being performed to drive the use of more accessible fluids^15,65,66^, practical diagnostic techniques remain difficult and are limited^13,14,67^. Our methods explore CWD testing options beyond current techniques by investigating biomarkers that can be performed in blood. We used plasma collected from CWD-infected Tg1536 mice along with controls and quantified the levels of three key biomarkers of neurodegeneration over the course of disease. NfL, GFAP, and t-Tau have been useful in diagnosing other neurodegenerative diseases^16,43–51,68–70^, yet have not been shown to be elevated in CWD plasma to date. Here, we demonstrated that all three of these biomarkers in plasma from CWD Tg1536 mice increased in concentration over time, correlated with disease stage, and were detectible at early stages of disease. Remarkably, all three biomarkers reached statistical significance at the early mid-timepoint of 132 dpi. The data suggests that we were able to capture the approximate time frame in which all plasma biomarkers reached significant elevation, as the 90 dpi samples had not all yet reached significance. By the next tested timepoint, 174 dpi, the statistical significance of all three biomarkers was strong, similar to levels observed at endpoint. While these biomarkers indicate neuronal damage, synaptic dysfunction, and astrogliosis are occurring^43,47,69,70^, they cannot be solely relied on to diagnose CWD due to their common presence in any condition related to CNS injury. Regardless, this study lays a crucial foundation demonstrating the utility of CWD plasma biomarkers.

### 4.4 Timeline of disease signs

Our CPP techniques detected CWD relatively early in the disease course, with both plasma biomarkers and behavioral signs emerging at early antemortem timepoints with most achieving significance in mid-timepoints. This innovative study began the groundwork of providing time-course investigations of blood-based diagnostic assays integrated with clinical signs of disease and neuropathology. An illustration of our CPP timeline is shown in **Figure 13**.

**Figure 13:**
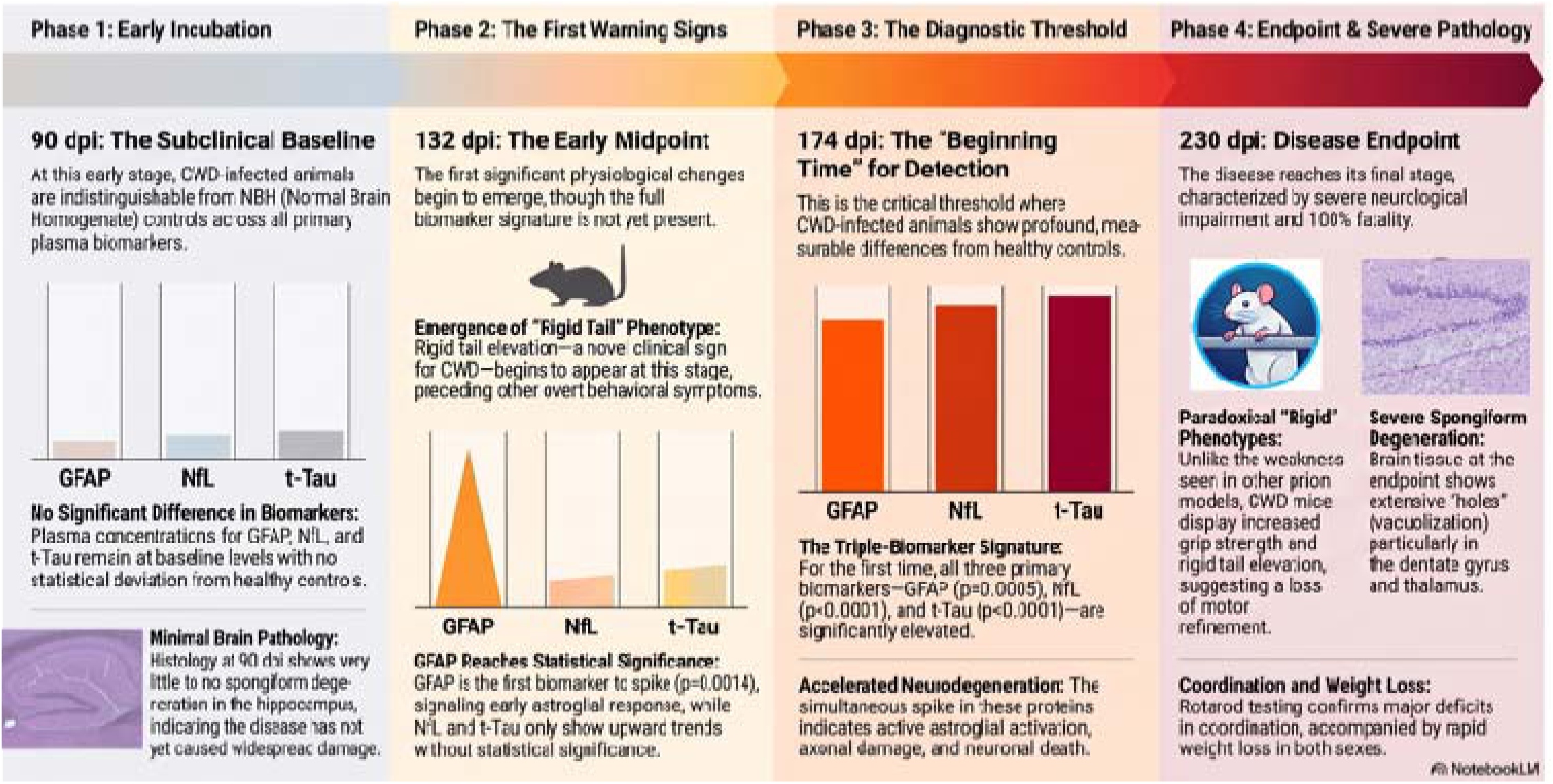
Tracking the Timeline of CWD Biomarker and Phenotype Progression. Figure generated by Notebook LM AI Platform.

## 5. Conclusion

Current CWD testing strategies are essential for monitoring disease spread in cervid populations; however, continued refinement of practical, minimally invasive testing approaches is needed to reduce environmental contamination, limit potential human exposure, and strengthen management strategies for both wild and farmed cervids. To our knowledge, plasma biomarkers of CWD infection in living animals have not previously been reported. In this study, we identified significantly elevated concentrations of NfL, GFAP, and t-Tau in CWD-infected transgenic mouse models, and these biomarker changes correlated with disease-associated phenotypes detected using our CPP methods. These findings support the potential utility of plasma biomarkers for live-animal assessment and future refinement of disease staging through less invasive sampling approaches, such as venipuncture. Importantly, preclinical biomarkers could provide a valuable tool for population-level disease management by enabling detection in asymptomatic animals and informing decisions related to animal movement, surveillance, and targeted intervention. Such applications may help reduce disease transmission through both direct animal-to-animal spread and environmental contamination.

## Ethical Statement

The research presented here was approved by the Weissman Hood Institute at Touro University; McLaughlin Research Institutional Animal Care and Use Committee under protocol number 2023-AG-100 and biosafety protocol number 2022-AG-IBC5.

## Acknowledgements

The authors thank Brent Race for contributing the CWD inoculum and Glenn Telling for providing the Tg1536 mice and thank you to both for being exceptional mentors. We thank Brenda Canine and Trisha Cox for use of the RT-QuIC plate reader and the analysis training. June Pounder’s contribution for mouse genotyping, study design, and inoculation assistance is greatly appreciated. Thanks to Bridget Gray for assistance with implementation of the grip strength assay and thoughtful discussions regarding the data analysis methods. We are grateful for Jon Buhrman’s training and assistance with the MSD platform.

Genotyping, colony management, rotarod, and DigiGait^TM^ behavioral assessments for the mouse models included in this study were provided by the McLaughlin Research Institute - Gene Editing and Mouse Models Assessment (GEMMA) Core Facility within the Center for Integrated Biomedical and Rural Health Research, RRID:SCR_027045, 1P20GM152335. The authors thank Rose Pitstick and Kaela Davy for exceptional care of the mice. Finally, we thank Renee Reijo Pera and the COBRE administrative core for inspiration, thoughtful discussions, and support with this study.

