## Supplementary Materials for "A New Frontier in CWD Detection: Antemortem Plasma Biomarkers and Behavioral Profiling in Transgenic Mouse Models"

| General Category | Assessment | Score Description | Score |
|---|---|---|---|
| Overall Condition/Appearance | Weight | Weight measured with scale | Weight in Grams to 0.1 |
| Response to Environment | Nesting | Nesting assessment of a nestlet | 0=>90% nestlet is torn apart and a nest has been built |
| | | | 1=75% nestlet is torn apart and an organized nest is made |
| | | | 2=50% nestlet is torn apart, may have an organized nest or nestlet is scattered all over cage |
| | | | 3=10% or less of nestlet has been touched. Edges may be frayed, but 90% nestlet intact |
| Neurological Signs | Tail elevation | Tail is elevated off floor when ambulating around cage, assessment is made after mouse is placed back into cage | 0=normal tail posture |
| | | | 1=mouse ambulates half cage with tail ~30% angle off horizontal |
| | | | 2=mouse ambulates full cage with tail 30-60% off horizontal |
| | | | 3=tail remains elevated as mouse is ambulating and is over 60% angle |
| | Clasp | Hind feet clasp together when mouse is picked up by tail | 0=no clasp |
| | | | 1=clasp briefly, <1 sec |
| | | | 2=clasp longer (>1 sec) and then separate |
| | | | 3=clasp and do not separate until mouse is placed back on floor |
| Motor Ability | Trunk curl | Assessment of trunk strength when picked up by tail to reach for food rack; abnormalities may be either weakness with no trunk stability or neurologic type curls to the side | 0=normal |
| | | | 1=mild abnormality |
| | | | 2=moderate abnormality |
| | | | 3=severely different than normal appearance |

Table S1: Detailed descriptions of the observational neurobehavioral profiling assessments under each general behavioral category and scoring parameters.

| Overall Condition/Appearance | Response to Environment | Neurological Signs | Motor Ability |
|---|---|---|---|
| Coat Condition | Home cage activity | Piloerection | Initial Activity |
| Nose bulge | Food grinding | Vocalization | Body Tone |
| Ear Position | | Righting Reflex | Limb Tone |
| Palpebral closure | | Ear Twitch | |
| Fighting | | Plastic tail | |
| Barbering | | Pelvic Posture | |
| Lordokyphosis | | Observational Gait | |
| Respiration Rate | | Ataxia | |
| | | Bar balance | |
| | | Forepaw reaching | |
| | | Tremor | |
| | | Myoclonis | |
| | | Bradykinesia | |
| | | Circling | |
| | | Paresis/Paralysis | |
| | | Head tilt | |
| | | Seizure Activity | |

Table S2: Observable traits that were not found to be significantly different between the CWD and NBH mice at any timepoint

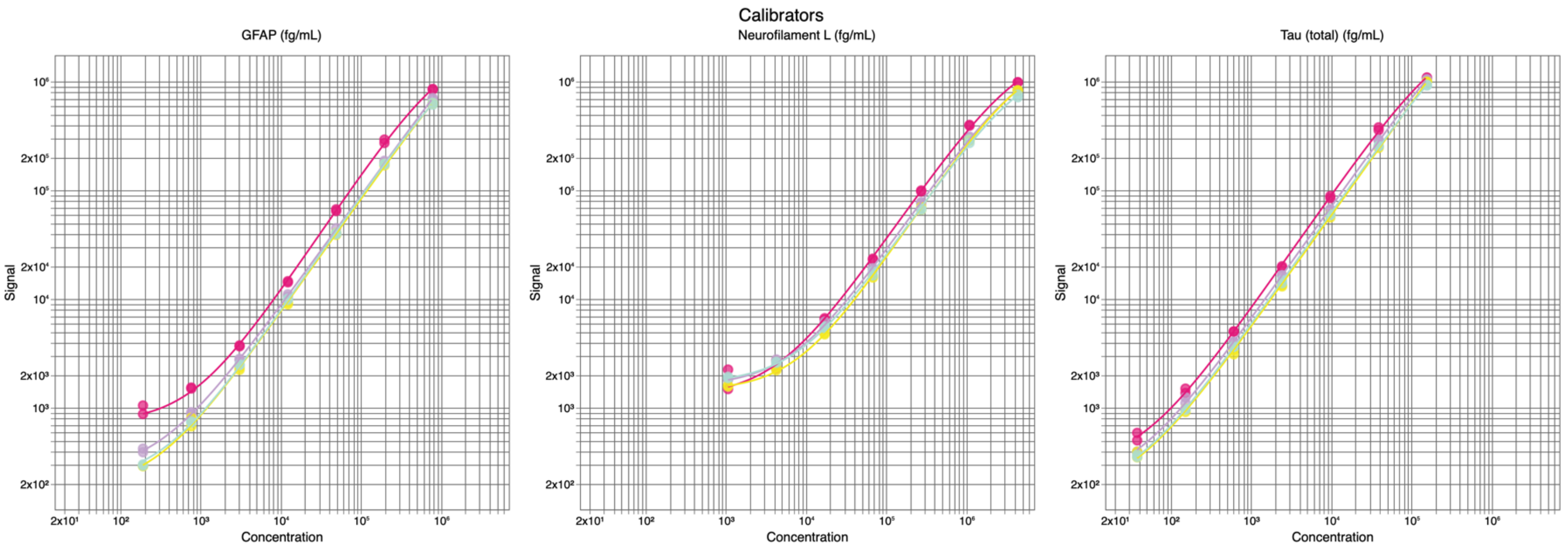

**Figure S1: Standard curves in plasma biomarker** MSD study used for Methodical Mind Enterprise data analysis in the early timepoint [blue], 132DPI [yellow], 174DPI [purple], and endpoint [pink].

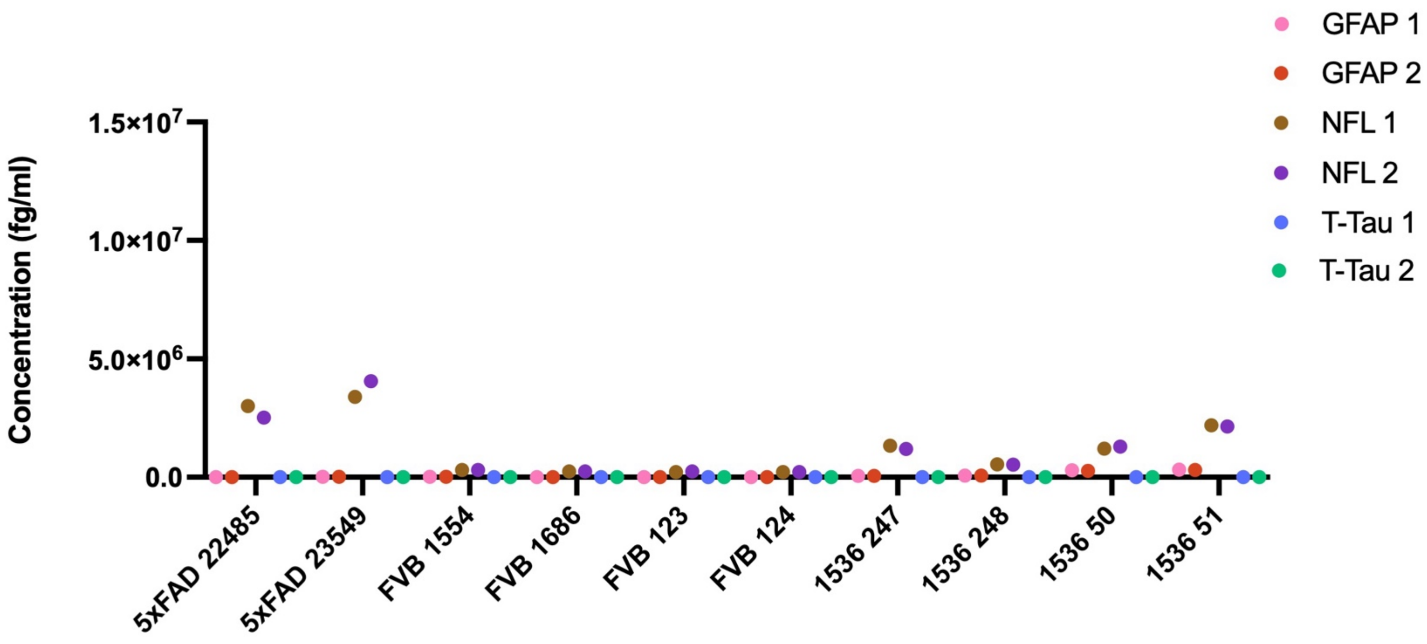

**Figure S2: Concentration of each replicate of GFAP, NFL, and Total-Tau** in control mice included on endpoint plate (230dpi). First and second replicate of GFAP per mouse is shown in pink and red, respectively. First and second replicate of NFL per mouse is shown in brown and purple, respectively. First and second replicate of Total-Tau per mouse is shown in blue and green, respectively.

A

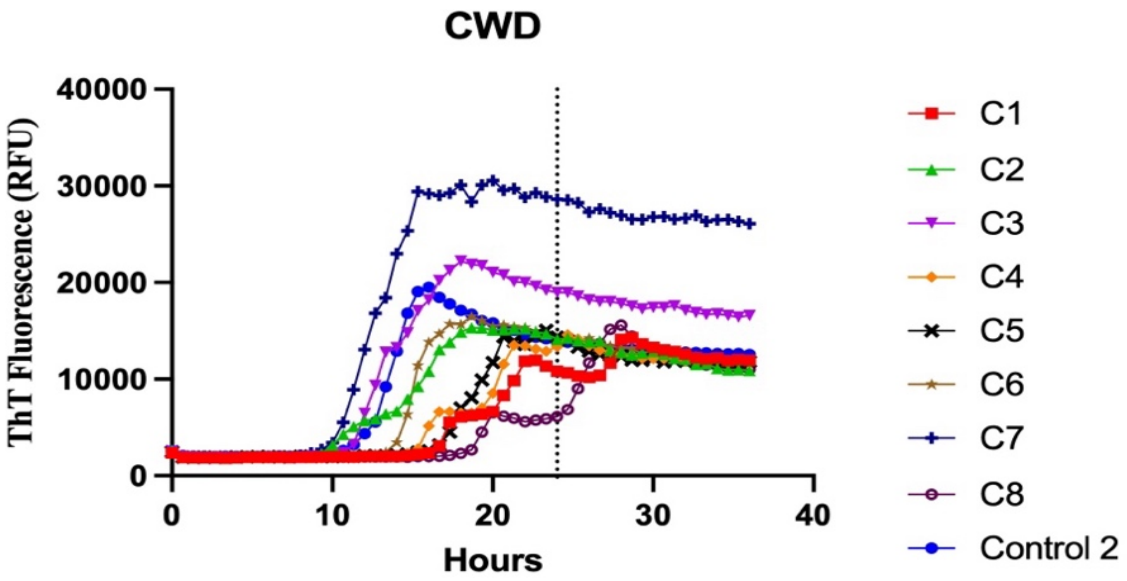

B

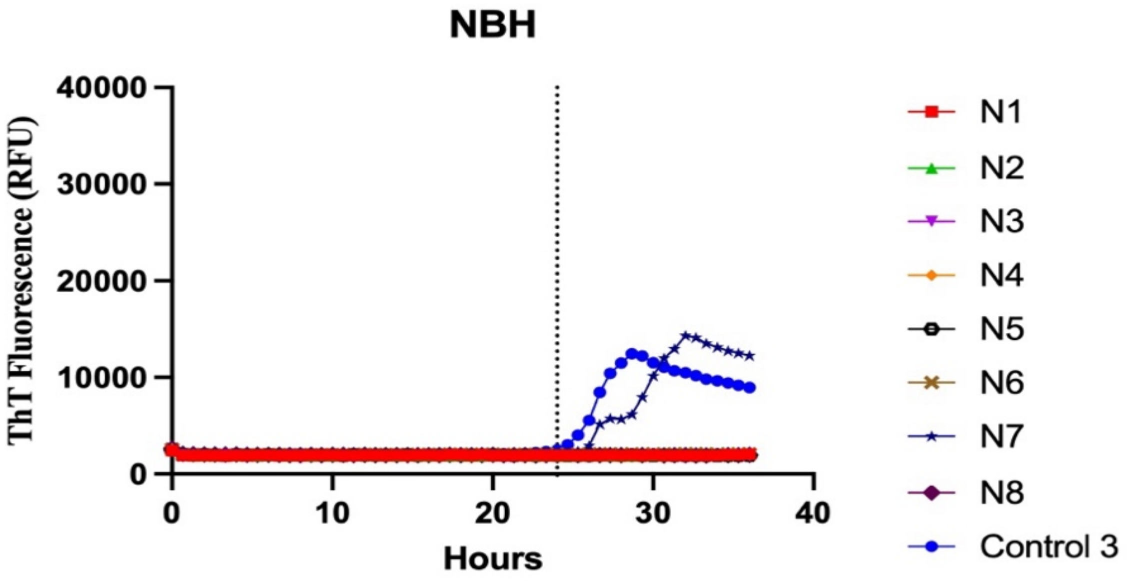

**Figure S3: Prion RT-QuIC kinetic data of samples as a mean of 4 replicates [A]** CWD inoculated Tg1536 brain homogenate and positive controls demonstrating seeding activity before cutoff at 24 hours. **[B]** NBH inoculated Tg1536 brain homogenates and negative controls demonstrating a lack of seeding activity before cutoff at 24 hours.
